# Molecular Disease Stages of Oligodendrocytic and Neuronal Tau Burden in Progressive Supranuclear Palsy

**DOI:** 10.64898/2026.08.03.742447

**Authors:** Nils Briel, Viktoria C. Ruf, Paul L.C. Feyen, Sigrun Roeber, Thomas Arzberger, Otto Windl, Tobias Weiss, Paolo Arosio, Günter U. Höglinger, Felix L. Struebing, Jochen Herms

## Abstract

**Background:** Progressive supranuclear palsy (PSP) is a primary tauopathy defined by the accumulation of 4R tau isoforms in neurons, oligodendrocytes and astrocytes. Despite evidence of genetic susceptibility operating through glial cell types, it remains poorly understood how cell type-specific epigenetic-transcriptional programs evolve with progression of tau pathology.

**Methods:** We conducted single-nucleus chromatin accessibility (snATACseq) and RNA sequencing (snRNAseq) on postmortem frontal cortex samples from PSP patients (n = 8) and matched controls (n = 8), yielding over 144,000 nuclei passing quality control. Tau pathology burden, including neurofibrillary tangles, coiled bodies, and tufted astrocytes, was quantified on AT8-immunostained sections from the same individuals. We integrated differential gene expression analysis, transcription factor motif enrichment, weighted gene co-expression network analysis, and pseudotime modeling anchored to cell type-specific tau pathology burden to delineate molecular pseudo-progression trajectories.

**Results:** In eight cell types, 20 subclasses, and 70 subclusters, PSP brains displayed a selective depletion of certain excitatory deep-layer neurons and oligodendrocyte subclusters, with relative preservation of inhibitory neurons and vascular cells. Genetic risk enrichment was localized to astrocytes and oligodendrocytes, whereas excitatory neurons exhibited the greatest transcriptional dysregulation. Oligodendrocyte pseudo-progression indicated a transition from homeostatic myelination programs (*MBP, MOBP*) through glucocorticoid-responsive stress (*FKBP5*, *ZBTB16*), to compensatory myelination (*PLP1, CNP*) and proteostasis stress (*UCHL1, CYRAB, CLU*). Neuronal pseudo- progression revealed early dysregulation of synaptic (*RORB2, NRG3, NPTX1*), microtubule dynamics (*KIF2C, RAB27B, TUBA/B*), and survival (*MEG3, FTX*) pathways, alongside a transient increase in neuron-glia interactions (*GRIP, CNTNAP4, ERBB4*), converging late on ribosomal translation and vesicular trafficking modules across all neuronal subtypes. Cross-modal integration with independent cerebrospinal fluid proteomics identified a concordant subset of glial reactivity, axonal injury, and synaptic markers jointly dysregulated in inhibitory neurons, oligodendrocytes, and excitatory deep-layer neurons.

**Conclusion:** PSP pathogenesis reflects a combination of glial genetic susceptibility and staged, cell type-specific transcriptional dysfunction. Oligodendrocytes transition from myelination-competent states to *FKBP5*-mediated stress states, while neurons show variably timed loss of synaptic excitability and survival programs, preceded by neuron-glia interactions and followed by convergent ribosomal-proteostatic failure. These cytopathology-anchored trajectories outline a potential pathophysiological sequence and may inform candidate selection for stage-specific therapeutic interventions in PSP.

## Background

Progressive Supranuclear Palsy (PSP) is a debilitating neurodegenerative disorder and the most common primary tauopathy. Clinically, PSP manifests as progressive motor, ocular-motor and cognitive impairment, significantly impacting quality of life and life expectancy [1]. The neuropathological hallmark of PSP is the accumulation of misfolded 4-repeat (4R) tau isoforms in neurons and glial cells, with a pattern of intercellular and cross-regional propagation [2–4].

While PSP shares some neuropathological features with Alzheimer’s disease (AD), such as tau aggregation in neurons, it is distinguished by the predominance of 4R tau isoforms and the involvement of astrocytes and oligodendrocytes in forming pathognomonic inclusions [5]. This highlights a unique role for glial cells in PSP pathogenesis. Cell type-specific vulnerabilities have been increasingly recognized as critical to understanding neurodegenerative diseases. For instance, cell types implicated in tauopathies include excitatory neurons broadly [6], specific excitatory neuron subtypes that are selectively depleted and preferentially harbor tau pathology [7], and astrocytes [8,9]. Astrocytes can accumulate tau and exhibit functional perturbations (e.g., altered synaptic-support- relevant programs and cellular homeostasis), reinforcing that glial pathology is not merely secondary to neuronal tau aggregation. In PSP, our prior work demonstrated that genetic risk loci identified through genome-wide association studies (GWAS)[10–12] are enriched in astrocytic epigenomic regions [13], unlike AD, where microglial genetic signatures predominate [14]. Furthermore, we previously identified transcription factor (TF) signatures associated with phosphorylated tau inclusions in astrocytes, which differed between PSP and corticobasal degeneration (CBD). Most recently, single-nucleus transcriptomic analysis of the PSP subthalamic nucleus revealed dysregulation of the integrated stress response across multiple vulnerable cell types, with eIF2α activation positively correlating with tau pathology burden [15]. However, the specific molecular cell states associated with the neuropathological progression of PSP remain poorly understood.

Currently available single-cell sequencing technologies enable unprecedented insights into cell type-specific transcriptional and epigenetic landscapes across disease states in primary and secondary tauopathies [6,13,16,17]. Moreover, the concept of multicellular “communities” has been introduced before [17], showing that coordinated co- variation of cell states across neurons and glial cell types tracks trajectories from healthy states to AD, emphasizing that disease-linked programs are emergent properties of interacting cell types rather than isolated populations. This framework motivates integrative analyses that relate coordinated regulatory programs to quantitative pathology measures, aligning with the rationale for combining single-nucleus molecular profiling with histopathological readouts.

In this study, we used single-nucleus RNA sequencing (snRNAseq) and single-nucleus ATAC sequencing (snATACseq) to generate a comprehensive multi-omic dataset from postmortem frontal cortex samples of PSP patients. By integrating these molecular profiles with quantitative neuropathological assessments of tau burden, we aimed to identify and understand the sequence of cell type-specific gene expression and epigenetic networks associated with disease progression. Understanding the molecular underpinnings of PSP is crucial for developing diagnostic biomarkers and disease-modifying therapies, as current diagnostic and treatment options remain limited [18,19].

## Methods and materials

### Ethics statement

All research at the Center for Neuropathology and Prion Research was conducted according to approved Brain Banking protocols (#345-13, Ludwig-Maximilians-University Munich) and in accordance with procedures approved by the Ethics Committee of the Ludwig-Maximilians-University Munich (#19-442).

### Neuropathologic examination and cohort characteristics

Human brain tissue was provided by the Neurobiobank München (NBM). PSP cases (n = 8) and controls (n = 8) were selected and matched to minimize co-pathology and differences in age, postmortem interval (PMI), and sex (**Table 1**). PSP was diagnosed based on established neuropathologic criteria NINDS [20,21]. Controls were negative for clinically documented neurologic or psychiatric disease, and for a significant neuropathology. Co-pathologies that disqualified for inclusion were AD pathology (Consortium to Establish a Registry for Alzheimer’s Disease, CERAD > 0), and the presence of Lewy body [22] or other α-synuclein inclusion pathology, TDP-43+ cytoplasmic inclusions, 3R tau (RD3), or 4R tau (RD4; in controls). In some cases, lesion types were identified outside the region of interest (**Table 1**).

**Table 1.** Cohort characteristics. <u>Abbreviations</u>: α-synuclein (α-syn), amyloid-β (Aβ; 4G8 clone), argyrophilic grain disease (AGD), Assay for Transposase-Accessible Chromatin using Sequencing (ATAC), Consortium to Establish a Registry for Alzheimer’s Disease (CERAD), fused in sarcoma (FUS), hours (h), Lewy bodies (LB), postmortem interval (PMI), years (y).

|  | Demographics |  |  |  |  | Histopathology |  |  |  |  |  |  |  |  |  | Omics data |
| --- | --- | --- | --- | --- | --- | --- | --- | --- | --- | --- | --- | --- | --- | --- | --- | --- |
| ID | Diagnosis | Age [y] | PMI [h] | Sex | APOE | Braak stage | Thal phase | CERAD | Aβ (4G8) | AGD | TDP-43 | FUS | LB | α-syn | RNA | ATAC |
| PSP1 | PSP | 68 | 38 | Male | - | 1 | 0 | 0 | no | no | no | no | no | no | yes | yes |
| PSP2 | PSP | 76 | 78 | Female | E3/E3 | 0 | 0 | 0 | no | no | no | no | no | no | yes | yes |
| PSP3 | PSP | 78 | 42 | Male | E3/E3 | 1 | 0 | 0 | yes | yes | no | no | no | no | yes | - |
| PSP4 | PSP | 68 | 7 | Female | E3/E3 | 0 | 0 | 0 | no | no | no | no | no | no | yes | yes |
| PSP5 | PSP | 59 | 28 | Male | E2/E3 | - | - | - | - | yes | no | - | no | no | - | yes |
| PSP6 | PSP | 67 | 32 | Female | E3/E3 | 3 | - | 0 | - | no | no | - | no | no | yes | - |
| PSP7 | PSP | 70 | 27 | Male | E3/E4 | 2 | 2 | 0 | yes | no | no | no | no | no | yes | yes |
| PSP8 | PSP | 80 | 16 | Female | E3/E3 | 2 | 1* | 0 | yes | no | no | - | no | no | yes | yes |
| C1 | control | 58 | 22 | Male | E2/E3 | 1 | 0 | 0 | no | no | yes | - | no | no | yes | yes |
| C2 | control | 87 | 27 | Male | E2/E3 | 2 | 1 | 0 | yes | yes | no | no | no | no | yes | yes |
| C3 | control | 85 | 20 | Female | E3/E3 | 1 | 0 | 0 | no | no | - | - | no | no | yes | yes |
| C4 | control | 64 | 33 | Female | E3/E3 | 1 | 0 | 0 | no | no | no | no | no | no | yes | yes |
| C5 | control | 53 | 29 | Male | E3/E3 | 1 | 2 | 0 | yes | no | no | no | no | no | yes | yes |
| C6 | control | 62 | 10-34 | Male | E3/E3 | 1 | 0 | 0 | no | no | no | no | no | no | yes | yes |
| C7 | control | 81 | 13 | Female | E3/E3 | 3 | 0 | 0 | no | no | no | no | no | no | yes | yes |
| C8 | control | 67 | 33 | Female | E3/E3 | 1 | 3 | 0 | no | no | no | no | no | no | yes | yes |

The region of interest was the medial and superior frontal gyrus at the level of the anterior striatum, corresponding to Brodmann areas 6/8/9. This cohort extends the snATACseq dataset acquired from cases PSP1-4 and control 1-5 that has been published previously [13].

#### Quantification of neuropathological traits

AT8 (Thermo Fisher Scientific, Waltham, MA, USA) immunohistochemical staining (diaminobenzidine) was carried out on 5-µm formalin-fixed paraffin-embedded sections from the contralateral middle frontal gyrus (MFG), corresponding to the cryo-frozen hemisphere used for single-nuclei sequencing, using a Ventana BenchMark Ultra stainer (Ventana Medical Systems, Roche Diagnostics, Tucson, AZ, USA) with an antibody dilution of 1:400.

Digitalization was performed on a Zeiss Axioscan 7 system with semi-automated scanning at 1x, 10x, 40x, and 100x magnification, following manual specimen boundary marking on scout images. Images were saved as .lcm files and loaded into ImageJ/Fiji for Linux arm64 at 40x magnification. Ten 800 µm × 800 µm regions of interest were randomly sampled across the slide (8 from cortex, 2 from white matter) to account for cortical cell overrepresentation in multi- omic analyses. Within these regions of interest, disease-defining AT8-positive intracellular inclusions were manually counted as classified as i) neurofibrillary tangles (NFT), ii) tufted astrocytes (TA), or oligodendrocytic coiled bodies (CB) (**Extended Data Fig. 6-7**). For neuropil threads, we used a semi-quantitative staging scheme ranging from 0 (no threads) to 4 (maximum threads). All continuous quantifications were rescaled to the 0/4 range for better comparability.

#### Nuclei isolation, partitioning and library preparation for snATACseq and snRNAseq

Nuclei isolation for was performed as previously described [13]. In brief, approximately 200 mg of cryopreserved brain tissue was gently homogenized on ice in 3.75 mL of chilled lysis buffer (10 mM Tris-HCl pH 8.0, 0.32 M sucrose, 0.34 mM DTT, 0.1 mM PMSF, 3 mM MgAc_2_, 5 mM CaCl_2_, 0.1 mM EDTA and 0.1% Igepal as well as RNAse inhibitor (Roche for RNAseq) using a Dounce homogenizer and transferred to 15 mL-ultracentrifugation tubes (Seton Open-top polyallomer centrifuge Tubes) and supplemented with an additional 2.25 mL of lysis buffer. Homogenates were carefully underlaid with 6.75 mL of sucrose buffer (10 mM Tris-HCl, pH 8.0, 1.8 M sucrose, 0.34 mM DTT, 0.1 mM PMSF, 3 mM MgAc_2_) and centrifuged for 1 h at 24,000 rpm at 4 °C. The resulting nuclei pellet was resuspended either in 1 × nuclei buffer (10x Genomics) for ATACseq or in 1× PBS supplemented with 1% BSA and RNAse inhibitor (0.2 U/µL) for RNAseq. Nuclei were subsequently quantified using a LunaTM fluorescence cell counter (Logos Biosystems, Anyang, South Korea) and the nuclei stock concentration was adjusted to target approximately 5,000-10,000 nuclei per case.

SnATACseq libraries were generated using the Chromium Next GEM Single Cell ATAC Reagent Kit v1.1 (10x Genomics, Pleasanton, CA, USA), according to the manufacturer’s instructions. Following transposition, nuclei were loaded onto a Chromium Next GEM Chip H for partitioning using the Chromium Controller. After DNA recovery and cleanup, sample indices were added and libraries were subjected to double-sided size selection. Final libraries were eluted in 20 µL Buffer EB (Qiagen, Hilden, Germany) and stored at −20 °C.

SnRNAseq libraries were generated using the Chromium Single Cell 3′ Reagent Kit v3.1 (10x Genomics) according to the manufacturer’s instructions. Nuclei were partitioned on a Chromium Controller, followed by reverse transcription to generate barcoded cDNA. After incubation, barcoded cDNA was recovered and purified using silane magnetic bead– based cleanup as specified by 10x Genomics. After cDNA amplification and SPRIselect-based cleanup, sequencing libraries were constructed according to the 10x Genomics protocol, including enzymatic fragmentation, end repair, adaptor ligation, and index PCR. Final libraries were eluted in 35 µL Buffer EB (Qiagen, Hilden, Germany) and stored at −20 °C.

Library quality and fragment size distributions for both snATACseq and snRNAseq libraries were assessed using an Agilent 2100 Bioanalyzer with a High Sensitivity DNA chip (Agilent, Santa Clara, CA, USA). Library concentrations were determined using the KAPA Library Quantification Kit for Illumina platforms (Roche Diagnostics, Mannheim, Germany). Sequencing was performed on an Illumina NovaSeq 6000 platform targeting ≥ 25,000 read pairs per targeted nucleus for snATACseq libraries and 20,000 read pairs per nucleus for snRNAseq libraries, respectively.

### Data processing and analysis

Statistical analyses were performed using R 4.4.3 in RStudio server on Arch Linux 6.13.8. The significance level for inferential tests was set at an p-value of ≤ 0.05 unless stated otherwise.

#### Preprocessing and quality control

Illumina binary base call files were converted to *fastq* using command line *bcl2fastq*, followed by demultiplexing and sequence alignment to the reference genome (GRCh38.p14) using the Cell Ranger ATAC v1.2.0 (*cellranger-atac count* for snATACseq), or the Cell Ranger v7.2 pipeline (*cellranger count* for snRNAseq).

For snRNAseq data, the expression matrix and corresponding metadata *singlecell.csv* files were loaded into R using *Seurat v5* [23]. The quality control (QC) during data loading excluded genes expressed in < 3 cells and excluded cells with expressed genes ≤ 200 or ≥ 6000, mitochondrial genes ≥ 5%, and hemoglobin genes > 0.5%. To identify doublet nuclei, we calculated doublet probability scores using *scDblFinder* v1.18 (input: raw count matrix; default arguments) (**Extended Data Fig. 1**).

The single-sample *Seurat* objects were merged into a joined object, followed by normalization using *SCTransform* v0.4.1, regressing out the number of RNA counts, age, and mitochondrial gene percentage. The first 30 principal components from principal component analysis (PCA) were used for dimensionality reduction and clustering. We then applied *Seurat*’s RPCA-integration method to *SCTransform* the normalized data matrix. Variable features were identified using the *FindVariableFeatures* function, selecting genes based on variance stabilizing transformation (VST). A first set of subclusters was identified with Idents set to “subcluster” cell identity using the *FindClusters* function with a resolution of 0.8. These subcluster annotations were overwritten later (see section “Cell Type Annotation and Subcluster Validation”). We then applied a further stage of QC, “QC1”, to filter out barcoded cells with high doublet probability and moderate to high mitochondrial read contamination. The filter cutpoints were defined as scDblFinder score ≥ 0.5 and precent mitochondrial reads ≥ 0.3, based on i) the histogram of doublet probability or mitochondrial reads scores, respectively, and ii) on visual examination of feature projections in Uniform Manifold Approximation and Projection (UMAP) (**Extended Data Fig. 2**). An additional QC filter step was applied after cell type annotation (see below). The final UMAP was calculated by using the first 30 dimensions of the RPCA integration. All downstream analyses were conducted with filtered data, unless stated otherwise.

For snATACseq data, sample-specific *bed* and corresponding metadata *singlecell.csv* files were loaded into R using *Signac* v1.14 [24]. Peaks were called in a sample-wise manner using *MACS2*, and a common peak set was defined by merging peaks from all samples. Peaks were filtered to retain those in the range of 20-10,000 bp. The first stage QC excluded nuclei with passed filters features < 500, peak region fragments < 2,000 or > 2,000, nucleosome signal > 4, TSS enrichment score < 2, reads in peaks < 10% (peak region fragments / passed filters) and blacklist ratio > 0.05 (**Extended Data Fig. 4**). To identify doublet nuclei, we used the *scDblFinder* package (input: raw count matrix; arguments: aggregateFeatures = TRUE, nfeatures = 25, processing = "normFeatures") and excluded cells with scDblFinder_score ≥ 0.5. Data were normalized using the term frequency-inverse document frequency method [24]. We identified the top features with a minimum cutoff of ’q0’ and performed singular value decomposition to reduce dimensionality. To correct for batch effects, we applied *harmony* integration [25] on the latent semantic indexing (LSI) reduction, grouping by cases. UMAPs were generated from uncorrected LSI dimensions 2-30 or harmony corrected low-dimensional representation. Gene activity scores were calculated with *Seurat’s* built-in function for the top 3,000 variable genes (VST) of the snRNAseq data. The gene activity matrix was normalized using *Signac’s* built-in function.

TF motif analysis was conducted using the “BSgenome.Hsapiens.UCSC.hg38” reference genome and the JASPAR2022 TF motif database [26]. C*hromVAR* v1.26 [27] was run to infer TF activity from the snATACseq data, including iterative matching of GC content and sampling average accessibility distribution of motif-containing peaks. Cell type-specific enrichment of GWAS risk variants was assessed using *gchromVAR* v.0.3.2 [28], an adaptation of *chromVAR* designed to integrate fine-mapped genetic variants with quantitative chromatin accessibility data. For each trait, GWAS summary statistics of PSP GWAS meta-analysis, AD GWAS, etc. were overlapped with snATACseq peaks, and variants within each peak were weighted by their posterior probabilities of being causal. Mann-Whitney rank-sum tests assessed whether trait-associated variants were preferentially enriched in cell type-specific versus bulk accessible chromatin regions, with Bonferroni correction applied across all tested combinations.

#### Cell type annotation and subcluster validation

Cell type annotation was performed using the *Azimuth* v0.5 reference mapping tool [29] built in *Seurat*, with “humancortexref" as reference and the "RNA" assay of the QC1-filtered *Seurat* snRNAseq object. The reference-mapping includes three levels of brain cell ontology, namely “predicted.celltype” (“neuronal” or “non-neuronal”), “predicted.subclass” (e.g.,”Pvalb neuron”), “predicted.cluster” (e.g.,”Pvalb neuron layer 2/3”), but lacks a cell type level less differentiated than “predicted.subclass” but more granular than the dichotomic “predicted.celltype” labels. Therefore, starting from subclasses we redefined the “celltype” into astrocytes (Astro), vascular niche (Vasc), excitatory upper layer neurons (Exc-ULN), excitatory deep layer neurons (Exc-DLN), inhibitory neurons (Inh-Neu), microglia and perivascular macrophages (Micro-PVM), oligodendrocytes (Oligo), and oligodendrocyte precursor cells (OPC). Hereafter, we use the abbreviated cell identity (e.g., ‘Astro’) to denote the specific cluster in this dataset, whereas the written term (e.g., ‘astrocyte’) refers to the broader cell type beyond this dataset. An additional filter step was applied after cell type annotation to exclude non-referenceable nuclei (predicted cell type annotation score < 0.4 adding 0.2 safety value, **Extended Data Fig. 2**). Cell type labels were validated by visual inspection of the canonical marker gene expression matrix (**Extended Data Fig. 3**).

For subclustering, a modified approach as described in [17] was performed on each single “predicted.subclass”, except for vascular niche cells which were combined into a single cluster due to low overall nuclei counts. Genes expressed in < 15 nuclei and non-coding or non-annotated genes were removed. The optimal cluster count and subcluster assignments were determined using the *scSHC* package v0.1 [30] which employs a model-based hypothesis testing framework for hierarchical clustering to overcome under- and over-clustering. Subclasses with > 250 cells were immediately subjected to normalization using *SCTransform* specifying 4,000 variable features (except for Vasc: 2,000 variable features), followed by PCA and UMAP. Low-abundant cell subclasses with 25-250 nuclei were merged with the most proximate subcluster in the subclass k-nearest-neighbors graph (n=10 nearest neighbors). Resulting subclusters with < 3 contributing samples (i.e., likely batch biases of single samples) were merged with the most proximate subcluster in the subclass k-nearest-neighbors graph. In these instances, reformed subclass-subcluster populations were subsequently processed similarly with *SCTransform*, PCA, and UMAP visualization.

For label transfer and co-embedding of both RNA and ATAC assay modalities, transfer anchors were first identified between the snRNAseq and snATACseq datasets using the Canonical Correlation Analysis in *Seurat*. RNA data were then transferred to the snATACseq object. The combined dataset was scaled and processed with PCA. Finally, UMAP was run on the combined dataset to visualize the co-embedded data.

#### Cell type composition analysis

Cell type composition was assessed using the *sccomp* R package v1.8 [31], which applies linear mixed-effects regression models to the count distributions of annotated cellular groups including cell type, predicted subclass, and subcluster. For each group, the distribution of cell type counts across samples was modeled with the composition as the response variable. Fixed effects included disease/control group labels (diagnosis), PMI, and age, while a random effect for individual sample identifier accounted for inter-individual variability and tissue sampling bias. Analyses were performed separately at each annotation level, with diagnosis, PMI, and age at death included as covariates in both the composition and variability models. The relationship between mean proportion and variability was modeled as bimodal. During model fitting, outlier samples were automatically detected and removed prior to inference. For each annotation level, group differences in cell composition were statistically tested using |log2FC| thresholds of 0.1 for cell type and subclass, and 0.2 for subclusters.

#### Supervised pseudotime analysis

To estimate a temporal order of gene expression and chromatin accessibility programs along a presumptive disease progression we used *psupertime* V0.2.6 [32]. This framework extends unsupervised pseudotime inference methods by the incorporation of time labels or disease stages as response variables in penalized ordinal logistic regression models, hence allowing for supervised pseudotime analysis. The cell subclass-specific SCT-transformed Seurat expression matrices and respective meta data were converted into SingleCellExperiment objects, which were eventually used as input for the *psupertime* function. We provided ordinal neuropathological trait quantification labels (i.e., per case NFT load) as response variables and set the selected genes to “all”, the assay type to “logcounts”, used built-in scaling and smoothing functions, and only included genes with a minimum expression of 0.1.

#### Differential expression analysis

We performed differential expression analysis to identify genes that were significantly altered between PSP and control samples across different cell types and cellular subpopulations. The analysis was conducted using the *Seurat v5* [23] and *DESeq2* v1.44 [33] packages in R. We followed the HBC Training documentation with an exception as described below (https://github.com/hbctraining/scRNA-seq_online/blob/master/lessons/pseudobulk_DESeq2_scrnaseq.md). Pseudobulk samples were created by aggregating snRNAseq data for each combination of donor, condition (PSP or control), and cell type using the *matrix.utils aggregate.Matrix* function. This approach helps to mitigate the impact of technical noise and takes into account the biological variability between samples. We then performed differential expression analysis at three levels of cellular resolution: (i) cell types, (ii) subclasses and (iii) subclusters.

For the broad cell type and subclass analyses, we used *DESeq2* to compare PSP samples to control samples. We applied a threshold for ≥ 1% of cells expressing a given gene, a false discovery rate (FDR) of ≤ 0.1 and an absolute log_2_ fold-change (logFC) threshold of 0.1. This relatively conservative threshold was chosen to balance between discovery potential and statistical robustness. For the more granular subcluster analysis where aggregation following multiply stratifying by group, sample, and cell identity result in small sample sizes for statistical comparison, we employed the non-pseudobulk *MAST* algorithm as built-in function in *Seurat v5*. Here, we used more stringent thresholds, with ≥ 25% of cell expressing a given gene, a FDR of ≤ 0.1 and a logFC threshold of ≥ 0.25. Downstream analyses included inspection of overall DEG profiles using PCAplot in *DESeq2*, using the top 10,000 features.

#### Cell type-resolved weighted gene co-expression network analysis

We employed high-dimensional Weighted Gene Co-expression Network Analysis (*hdWGCNA,* v0.3.03) [34] to identify and characterize gene co-expression networks in the snRNAseq data.

We first prepared the Seurat object for *hdWGCNA* analysis using the *SetupForWGCNA* function. We selected genes based on single gene expression in a fraction of ≥ 0.05 nuclei. Metacells were generated grouping nuclei by their cell type and sample ID, using 25 nearest neighbors (k = 25) for metacell construction, with a ≤ 10 shared neighbors allowed between metacells. Network construction and module detection was performed by iterating through each cell type following the *hdWGCNA’s* native functions: 1) Soft power selection: the optimal soft-thresholding power was determined by aiming for a scale-free topology fit index (R^2^) > 0.8, selecting the lowest power that met this criterion. 2) Network construction: the co-expression network was constructed by applying the optimal soft power, and setting a module size of ≥ 50 genes to focus on larger, more robust gene modules; the topological overlap matrix was computed and used for module detection. 3) Module eigengene calculation: module eigengenes (i.e. 1^st^ principal component of each module’s expression profile), were computed after PCA, while regressing out the effects of age and PMI to account for potential confounding factors. 4) Module connectivity analysis: intramodular connectivity and module membership of each gene were calculated, which allowed for identifying hub genes (i.e., highly connected genes within their respective modules with a presumably essential function within that module).

Differential Module Eigengene (DME) analysis of PSP vs. control groups was performed using the *FindDMEs* function. We visualized module membership with gene dendrograms. We used the *ComplexHeatmap* package 2.20 [35] to generate a heatmap displaying DME average log2FC values between PSP and control samples across cell type sub-identities, with statistical comparisons between the nuclei MEs of a given sub-identity and the remaining nuclei as assessed by Wilcoxon rank-sum tests and Bonferroni correction. The heatmaps incorporate common annotations including module colors, subcluster abundance changes in PSP (if applicable), top hub genes, and enriched GO terms.

#### Gene ontology analyses

Unless stated otherwise, we used Gene Ontology (GO) enrichment analysis for functional annotation of gene sets resulting from differential expression analysis, hdWGCNA module hub and member genes and pseudotime- associated genes. Queries were made with *enrichGO* function in *clusterProfiler* v4.12.6 [36] referencing to the *org.Hs.eg.db* database v3.19.1. In all GO analyses the BH method was used to control the false discovery rate, and the minimum or maximum gene set sizes were set as 10 or 500 genes, respectively. For DEGs, GO analysis was conducted on each cell type, subclass or subcluster separately, querying up- or down-regulated genes, and setting a p-value cutoff < 0.05, q-value cutoff < 0.1, and all genes detected in the RNA dataset as background. For hdWGCNA modules, hub genes and members with a module eigenvalue of > 0.05 were subjected to GO analysis in a module- wise fashion applying a p-value cutoff of 0.05, q-value cutoff of 0.1, and all genes included in the co-expression network construction process as background. A maximum of 5’000 query genes was permitted for module genes.

#### Cross-modal integration with published cerebrospinal fluid proteomics

To determine whether cell type-specific transcriptional alterations in PSP frontal cortex are reflected in the cerebrospinal fluid (CSF) proteome, we integrated our snRNAseq DEG results with a published CSF proteomics dataset by Jang et al. [37] that includes PSP and healthy control (HC) samples. Parkinson’s disease and quality control specimens were excluded. Differential protein abundance between PSP and HC was estimated using the *limma* package v.3.66 [38] with empirical Bayes moderation, incorporating the processing batch as a covariate in the design matrix; the group coefficient was used downstream. For cross-modal concordance analysis, snRNAseq DEGs (this study, all cell types; **Extended Data Table 3**) were intersected with CSF-quantified proteins from Jang et al [37] by gene symbol. For each gene-cell type pair present in both datasets, the direction of regulation was classified as concordant upregulated (log2FC_snRNAseq_ > 0 and log2FC_proteomics_ > 0), concordant downregulated (both < 0), or discordant. A mean fold-change was computed as the arithmetic mean of the log2FC_snRNAseq_ and log2FC_proteomics_. Pearson correlation between the two fold-change vectors was calculated per cell type (and all CSF proteomics candidates). Concordant gene-cell type pairs were retained at thresholds of p_snRNAseq_ < 0.05 and of p_proteomics_ < 0.05 and |mean log2FC| > 0.5 for visualization.

## Data availability

The datasets analyzed during the current study can be made available upon request. The complete R scripts detailing the analysis pipeline are available under https://github.com/nes-b/PSP_multiomics.

## Results

### Cell type-specific transcriptome regulation in PSP

We created a high-quality multi-omic dataset of the PSP frontal cortex using case-paired single-nucleus ATAC- and RNA-seq to assess tauopathy-associated changes in transcriptome and chromatin accessibility jointly (**Fig. 1a,b, Extended Data Fig. 5**). The cohort consisted of eight PSP and eight age-, sex- and PMI-matched control cases, whereas both modalities were available for 13/16 donors (**Table 1**, **Extended Data Table 1**). We retrieved a total of 144‘296 nuclei after 1^st^ stage QC, with 55’413 high-quality nuclei from snATACseq and 88’883 high-quality nuclei from snRNAseq. These nuclei could be reattributed to three cell classes, encompassing eight major cell types and 19 subclasses (**Fig. 1c,d**). However, 8.98-12.23% of RNA nuclei were not sufficiently attributable to a reference subclass (i.e., had low prediction certainty, <u>see Methods</u>), and thus were excluded from downstream analyses leaving 138’012 nuclei (52’689 ATAC; 85’323 RNA). After integration with the RNA dataset, the ATAC nuclei were annotated accordingly. The final integrated snATAC/RNAseq dataset comprised a median of 3’889 (ATAC) and 5’788 (RNA) nuclei per control case, and 2’230 (ATAC) and 4’755 (RNA) nuclei per PSP case (**Extended Data Table 1**). The median count of ATAC peaks was slightly higher in controls (median [interquartile range]: 8‘302 [6’704, 9’098]) than in PSP cases (7’833 [7’141, 8’246]). Meanwhile, RNA features were balanced between both groups, with 5’510 [4’259, 6’352] in controls and 5’526 [3’527, 6’450] in PSP.

**Figure 1.**
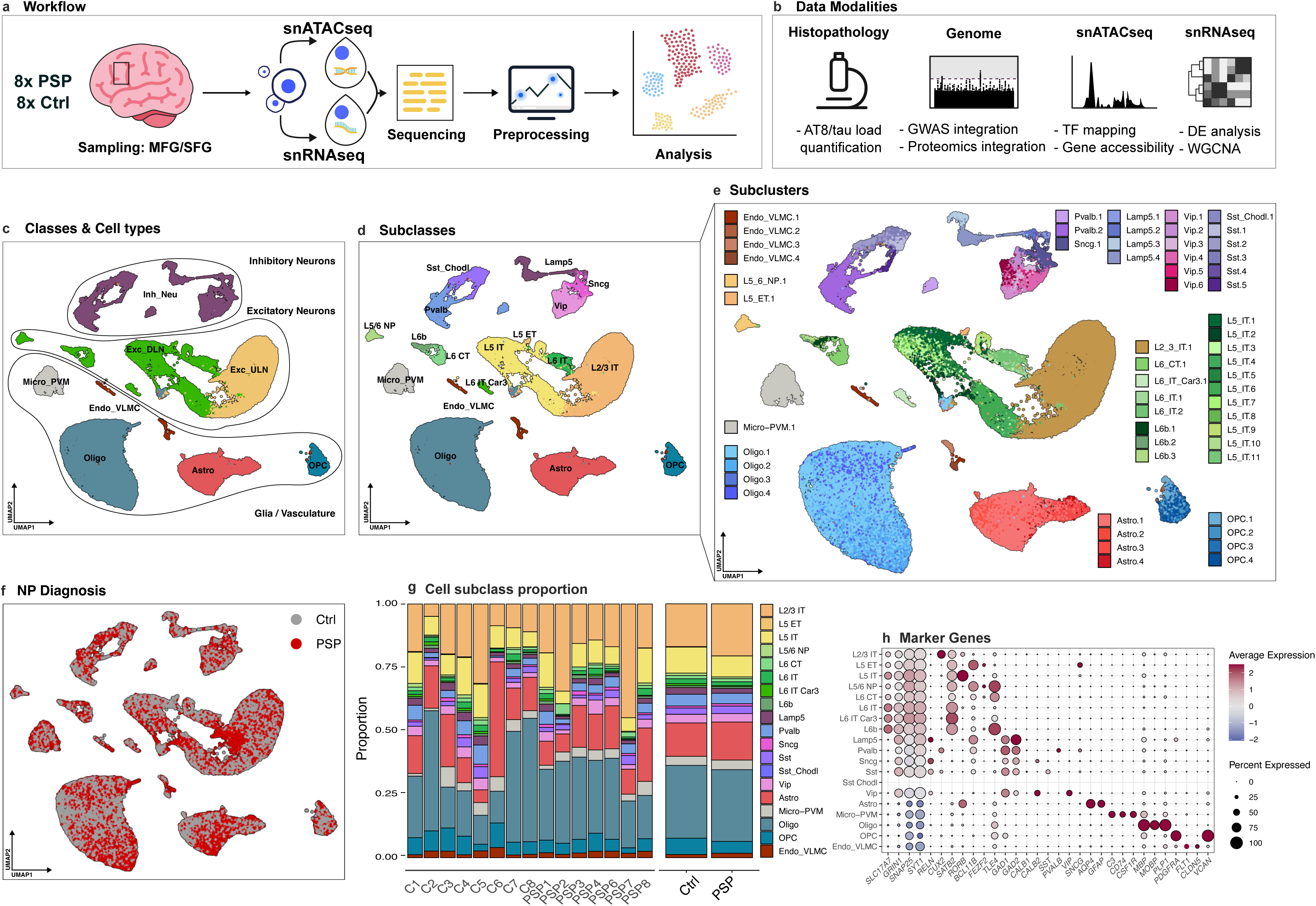
Workflow and single-nucleus profiling of PSP cortex. **(a)** Experimental workflow: Brain samples from 8 individuals with PSP and 8 controls were collected from the MFG and SFG. snATACseq and snRNAseq were performed, followed by preprocessing and integrative analysis. **(b)** Multi- modal data integration included histopathology (AT8/tau load quantification), genomic analysis (GWAS integration, TF mapping, gene accessibility), and transcriptomic analysis (DE analysis, WGCNA). **(c)** UMAP visualization of major cell classes identified in the dataset, including neurons, oligodendrocytes, astrocytes, microglia, endothelial cells, and vascular cells. **(d)** Subclasses within major cell types are further resolved into distinct populations such as inhibitory neurons, excitatory neurons, OPCs, astrocyte subtypes, and microglia subtypes. **(e)** Subclusters within subclasses reveal finer cellular diversity, including specific neuronal subtypes (e.g., L2/3 IT neurons), and glial subtypes (numbered consecutively). **(f)** UMAP projections highlight differences in cell populations between PSP and control samples. **(g)** Proportional representation of cell subclasses across PSP and control samples demonstrates shifts in cellular composition associated with disease state. **(h)** Dot plot showing marker gene expression across identified subclasses, highlighting genes characteristic of specific cell populations. <u>Abbreviations</u>: astrocytes (Astro), control (Ctrl), differential expression (DE), endothelial cells and vascular leptomeningeal cells (Endo_VLMC), excitatory deep-layer neurons (Exc_DLN), excitatory upper-layer neurons (Exc_ULN), genome-wide association study (GWAS), Lamp5-expressing interneurons (Lamp5), layer 2/3 intratelencephalic neurons (L2/3 IT), layer 5 extratelencephalic neurons (L5 ET), layer 5 intratelencephalic neurons (L5 IT), layer 5/6 near-projecting neurons (L5/6 NP), layer 6 corticothalamic neurons (L6 CT), layer 6 intratelencephalic neurons (L6 IT), microglia and perivascular macrophages (Micro_PVM), middle frontal gyrus (MFG), oligodendrocyte precursor cells (OPCs), oligodendrocytes (Oligo), parvalbumin-expressing interneurons (Pvalb), somatostatin-expressing interneurons (Sst), somatostatin/Chodl-expressing interneurons (Sst_Chodl), superior frontal gyrus (SFG), transcription factor (TF), uniform manifold approximation and projection (UMAP), vasoactive intestinal peptide-expressing interneurons (Vip), weighted gene co-expression network analysis (WGCNA).

Final annotations were consistent with canonical marker gene expression (**Fig. 1c-e,h**) and the nuclei showed sound integration across diagnosis groups (**Fig. 1f**), although there was a modest insignificant shift from neuronal towards glial (mainly oligodendro- and astrocytic) nuclei in PSP cases (**Fig. 1g, Extended Data Fig. 3c,d**, **7c**). Notably, several subclusters within the glial and excitatory neuronal compartments were either enriched or depleted in PSP, while inhibitory neuron subclusters (i.e., Pvalb, Lamp5 and Sst+) were present at smaller numbers in PSP (all FDR < 0.05, **Extended Data Fig. 7d**). This finding could indicate disease-specific cell type vulnerability and loss of selected neuronal cells, and vulnerability or cell state shifts in glia, e.g. from homeostatic towards reactive.

To investigate molecular changes over disease progression, we used the tau protein load as a surrogate marker. Based on AT8-stained histological slices from the frontal cortex of all PSP cases, we applied a semi-quantitative grading scheme for NFT, CB and TA to assess tau cytopathology burden and infer a pseudo-trajectory of tau accumulation. (<u>see Methods</u>, **Extended Data Fig. 7a**). Interestingly, with increasing NFT burden there were nominally relatively fewer layer 2/3 intratelencephalic (L2/3 IT), and with increasing CB burden relatively fewer layer 5 and 6 IT neurons, potentially reflecting layer-specific vulnerabilities with upper-cortical vs. deep-cortical/white matter tau pathology in PSP (**Extended Data Fig. 7b**). Furthermore, correlation analyses examined diagnostic Braak *neuroanatomical* staging (tau distribution across different brain areas) with *cytopathological* grading (local histological tau burden). These analyses revealed significant positive correlations between CB load and Braak staging of NFT. However, there was no correlation between CB and *cytopathological* NFT loads (**Extended Data Fig. 6a**). This suggests potential disease interactions of PSP-type and AD-type pathologies on large-scale neuronal networks.

### Cell type-specific epigenetic and transcriptional architecture of PSP

To replicate our previous findings that known GWAS risk variants for PSP and frontotemporal dementia were enriched in astrocytic ATAC peaks in a combined PSP+CBD cohort, we applied GWAS integration with snATACseq analysis using updated genetic database information [39] within an extended cohort of 7 PSP cases.

This revealed nominally significant results, none of which withheld multiple testing correction (**Fig. 2a**). Nonetheless, astrocytes demonstrated highest nominal enrichment of PSP genetic risk (p = 0.0329) followed by Oligo (p = 0.0442), supporting the notion that genetic predisposition to PSP operates primarily through glial dysfunction. Notably, across further disease-specific GWAS risk variants, there were enrichments of AD (p = 0.00369) and CBD (p = 0.00944) variants in the Exc-ULN population. Parkinson’s disease risk variants were enriched in Micro-PVM (p = 0.0186) and Exc-DLN (p = 0.0338; mainly deep-layer subclasses).

**Figure 2.**
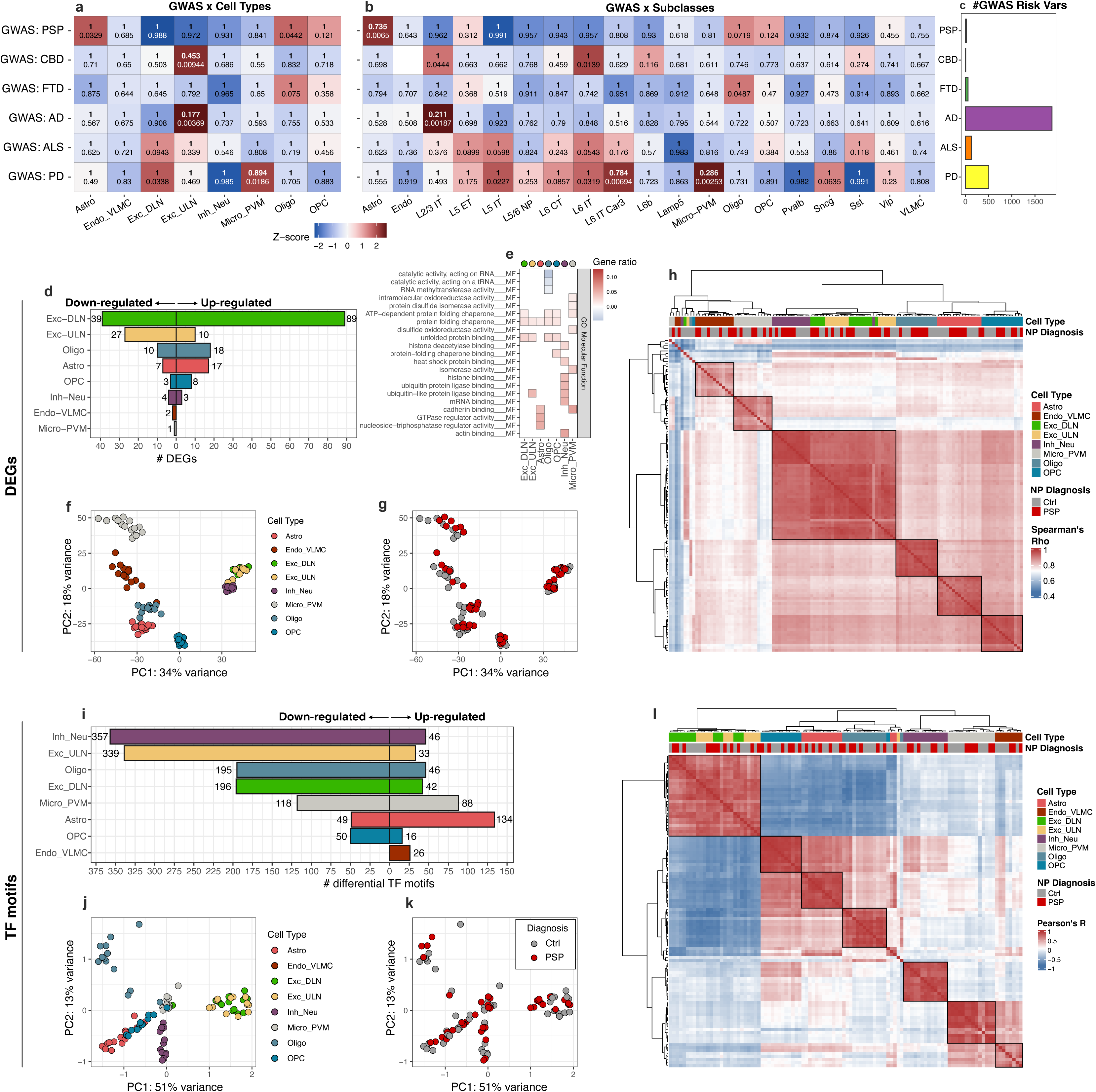
Multi-modal analysis of PSP-associated molecular changes across cell types and subclasses. **(a)** Heatmap showing enrichment of GWAS risk variants across major cell types computed with gchromVAR, with Z- scores as color fill reflecting the association between disease-associated risk variants and specific cell populations. Bonferroni-corrected p-values from Mann-Whitney rank-sum tests indicated as bold labels, nominal p-values in normal font below. **(b)** Heatmap showing enrichment of GWAS risk variants across cell subclasses. Plot structure same as (a). **(c)** Bar plot of absolute number of risk variants per disease annotated in the GWAS catalogue. **(d)** Bar plot of absolute numbers of DEGs by cell type and direction of change, underlying cut-off: adjusted p-value ≤ 0.1. **(e)** Heatmap of gene ratio values from GO and KEGG pathway gene set enrichment analysis using enrichGO, split by cell types on columns and top selected molecular function (MF) terms on rows. **(f, g)** PCA plots of DEGs by case and cell type grouping (pseudo-bulk), color indicates cell type **(e)** or group **(f)**. **(h)** Correlation heatmap of gene expression of DEGs by case and cell type grouping (pseudo-bulk), indicating Spearman’s Rho, ordered by hierarchical clustering. **(i)** Bar plot of absolute numbers of differential TF motifs by cell type and direction of change, underlying cut-off: adjusted p-value ≤ 0.1. **(j-k)** PCA plots of differential TF motifs by case and cell type grouping (pseudo-bulk), color indicates cell type **(j)** or group **(g)**. **(l)** Correlation heatmap of motif enrichment of differential TF by case and cell type grouping (pseudo-bulk), indicating Pearson’s R, ordered by hierarchical clustering. <u>Abbreviations</u>: Alzheimer disease (AD), amyotrophic lateral sclerosis (ALS), astrocytes (Astro), control (Ctrl), corticobasal degeneration (CBD), differentially expressed genes (DEGs), endothelial cells and vascular leptomeningeal cells (Endo-VLMC), excitatory deep-layer neurons (Exc-DLN), excitatory upper-layer neurons (Exc- ULN), frontotemporal dementia (FTD), gene ontology (GO), genome-wide association study (GWAS), inhibitory neurons (Inh-Neu), Kyoto Encyclopedia of Genes and Genomes (KEGG), microglia and perivascular macrophages (Micro-PVM), oligodendrocyte precursor cells (OPCs), oligodendrocytes (Oligo), principal component analysis (PCA), Parkinson disease (PD), transcription factor (TF).

The pseudo-bulk DEG analysis at the level of all cell types identified 479 DEGs (FDR < 0.1), with excitatory neurons exhibiting the greatest DEG burden, followed by Astro and Oligo (**Fig. 2d**). There was a good separation of cell type identities in case-wise collapsed transcriptomes in PCA consistent with moderate to high between-class correlations in the gene expression correlation matrix. When visualized by diagnosis labels, the PCA revealed most pronounced separation within glial subsets (**Fig. 2f-h**). Gene expression dysregulation and associated GO terms manifested in distinct patterns (**Fig. 2e, Extended Data T1, Extended Data Fig. 8**), where excitatory neurons displayed upregulation of proteostasis pathway-related genes (*HSP90AA1*, *TFEB*), and downregulation of synaptic-neuronal integrity markers (*CNTN6*, *BDNF)*. Astro activated genes related to unfolded protein response (*UBASH3B)* – as all other cell types – and increased extracellular signalling reception (*ICAM1, PLXND1, DUSP3)*. Oligo showed depletion of RNA metabolism and vesicle trafficking (long non-coding RNAs, *NAV3*), countered by induction of chaperones (*ZBTB16*, *FKBP5*). Micro-PVM gene expression was relatively inert and restricted to elevated chaperone, lysosome- endo-exocytosis and migration pathway activity (*RAB27A*, *FLOT2*).

The pseudo-bulk differential transcription factor motif enrichment analysis at cell type level identified 1’686 TFs (FDR < 0.1), with inhibitory neurons exhibiting the greatest differential TF burden, followed by excitatory neurons (**Fig. 2i**). There was less clear separation of case-wise collapsed cell type identities in the PCA, mirrored by high within- but low between-cell class correlation in the expression correlation matrix; diagnosis-based differentiation in PCA was moderate (**Fig. 2i,j**). Exc-DLN showed a marked reduction of both ROR- (RORB, RORA) and ETS-family (ELK1, ETS1) motifs, two essential neuronal gene regulator networks (**Extended Data Fig. 9**). Concurrently, these neurons gained STAT1 and STAT2 motif accessibility, indicating increased sensitivity to interferon-responsive signaling. In Astro, the TF regulation shifted towards a reactive phenotype: a gain of FOS and JUN motifs (immune activation and stress response) was paralleled by a decline in FOXO-family motifs, which have pleiotropic homeostatic functions [40]. Oligo displayed elevated accessibility at gluco-/mineralocorticoid receptors NR3C1/2 complementing alterations of the glucocorticoid-ZBTB16-FKBP5 feedback loop [41] and thus highlighting a stress-activated transcriptional state. Meanwhile, Micro-PVM exhibited a shift toward a pro-inflammatory phenotype, defined by increased motif accessibility for IRF4/8/9 and STAT1/2.

Together, the integrated epigenetic and transcriptomic data suggest that PSP pathogenesis involves glial-weighted genetic susceptibility, but broad cell type involvement with identity-specific dysregulation of proteostasis, synaptic integrity and stress-response pathways. TF regulation and transcriptional dysregulation showed a cell-type-specific dissociation.

### Oligodendrocytes shift from myelination to *FKBP5*-associated stress programs along tau progression

In PSP, oligodendrocytes harbour the hallmark cytopathology of CB which are 4R tau inclusions forming wire-like tangle arrangements in the cytoplasm. Based on the observations that Oligo subclusters exhibited the highest variability in cell abundance among all evaluated cell types (**Extended Data Fig. 7d**), that PSP GWAS risk variants localized to differentially accessible regulatory regions in PSP Oligo (**Fig. 2a**), and that PSP Oligo differentially express genes associated with cellular stress (such as *NAV3*, *ZBTB16*, and *FKBP5*; **Fig. 2c**), we hypothesized that oligodendrocytes transition into this stress-induced state in a gradual manner. To track transcriptional changes during the development of oligodendrocytic tau pathology, we modelled disease progression in Oligo using CB burden as a surrogate for cytopathology-specific disease progression (i.e., ‘pseudo-progression’, **Fig. 3a**, <u>see Methods</u>).

**Figure 3.**
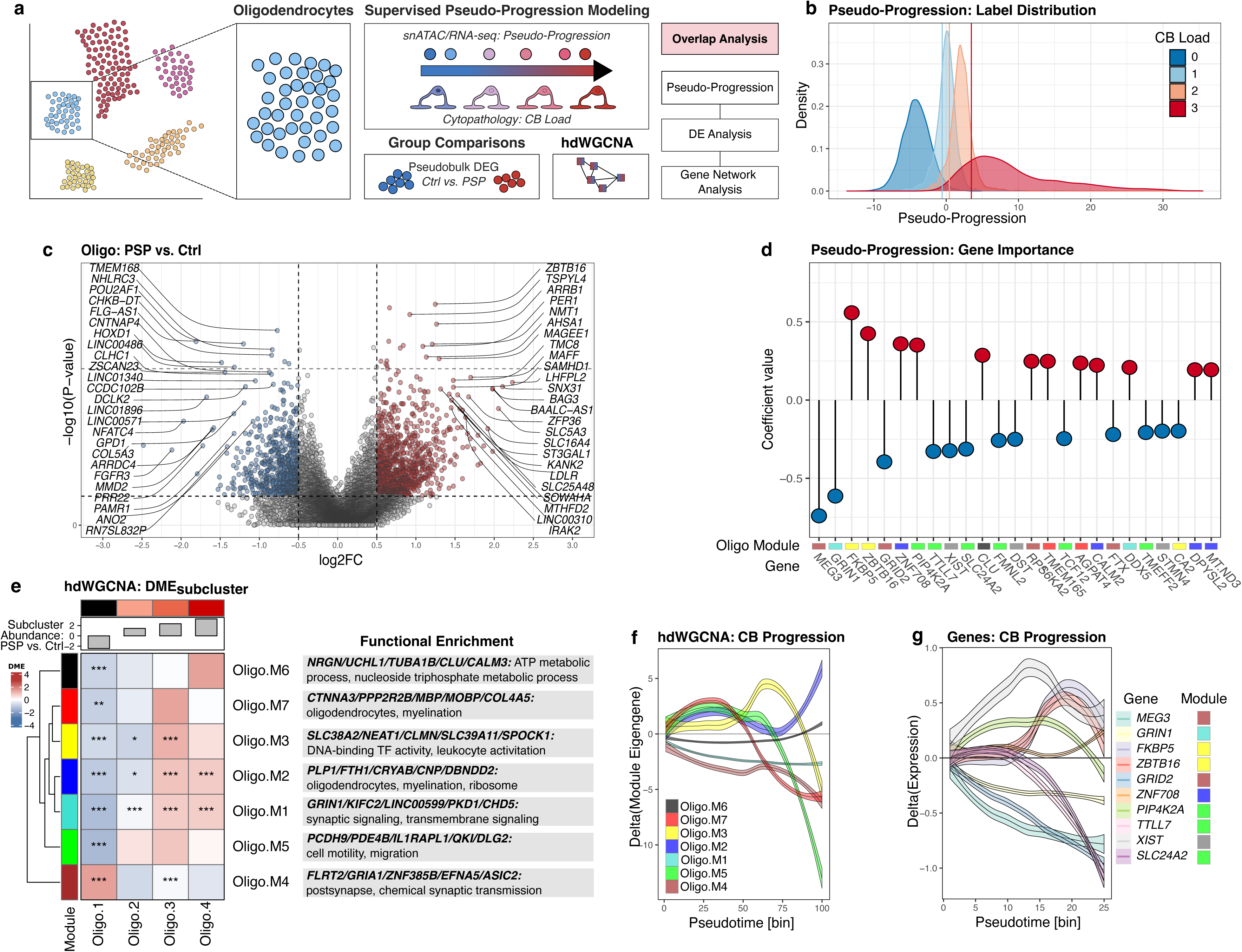
Oligodendrocyte transcriptional reprogramming along tau-pathology trajectories in PSP. **(a)** Analytical framework concept starting with the selection of Oligo subpopulation and association testing of omic data with increasing cytopathology CB load, i.e. modelling of ‘pseudo-progression’. In parallel, group comparisons through pseudo-bulk DEG and cell type-resolved WGCNA analyses were conducted. **(b)** Density plot of CB load labels over estimated pseudo-progression. **(c)** Volcano plot from pseudo-bulk DEG testing comparing PSP and control snRNAseq Oligo cell type data. Top 25 up- and downregulated genes are labelled. **(d)** Lollipop plots of Oligo gene expression pseudo-progression associations. Coefficient values of single genes are indicated, as provided from the final CB model fit in the Oligo cell type data. Color tiles at the bottom indicate hdWGCNA module assignment (see (e)). **(e)** Heatmap displaying DME of PSP and control snRNAseq Oligo cell type data, stratified by Oligo subcluster identity and seven distinct hdWGCNA modules. Top panel indicates subcluster abundance in PSP (positive) vs. control (negative values). Right-hand panels indicate GO enrichment of each module; top 5 hub genes and top functional enrichment terms are indicated. **(f)** Loess fit of hdWGCNA module eigengene values over CB pseudotime (i.e., pseudo-progression). Delta values are provided relative to the baseline. Color indicates Oligo hdWGCNA modules. **(g)** Loess fit of top 10 pseudo-progression-associated genes over CB pseudotime. Delta values are provided relative to the baseline. Left color legend column indicates genes, right color legend column indicates the respective Oligo hdWGCNA modules. <u>Abbreviations</u>: coiled body (CB), differential module eigengene (DME), high-dimensional weighted gene co- expression network analysis (hdWGCNA), log2-fold-change (log2FC), oligodendrocyte (Oligo).

In unsupervised scSHC subclustering [30] we identified four replicable Oligo subclusters with distinct transcriptional signatures and differential associations with CB burden (**Fig. 3b,e**). Subcluster Oligo.2, characterized by low myelination-related transcriptional programs, together with Oligo.3 and Oligo.4, both marked by enrichment of mature oligodendrocyte markers, were significantly enriched in PSP, whereas Oligo.1, defined by intercellular signalling and low myelination programs, was relatively depleted (**Extended Data Fig. 7d**). The pseudo-progression model trained on semi-quantitative CB load demonstrated a reasonable prediction accuracy (nucleus-to-CB-stage assignment: training set 74%, test set 73%), indicating that Oligo transcriptional profiles systematically vary as a function of tau cytopathology severity, while allowing overlapping CB-load labels to capture gradual within-case variation along the pseudo-progression scale (**Fig. 3c,d**). Pseudo-progression-associated gene expression patterns revealed a progressive shift from homeostatic myelination programs towards stress response activation. Among individual genes, *FKBP5* showed the strongest positive correlation with CB burden (β = 0.51, p = 0.001), aligning with its known role as a co-chaperone in tau oligomerization and stress-induced transcriptional reprogramming.

Gene network analysis using hdWGCNA identified seven Oligo-specific modules, which varied nonlinearly with increasing CB burden (**Fig. 3e,f, Extended Data Fig. 10**). Modules M1 (turquoise) and M4 (brown), both associated with synaptic function and cell adhesion, declined early and slowly with increasing pseudo-progression. Similarly, Module M6 (black), driven by proteostasis and microtubule hub genes (*UCHL1, CLU, TUBA1B*), displayed a slow dynamic but increased with late pseudo-progression.

Both modules M5 (green), enriched for genes involved in cell motility and migration, and a myelination-associated module M7 (red), exhibited an early rise followed by a progressive downregulation during mid-scale pseudo- progression. Conversely, module M3 (yellow), dominated by stress-response genes (*SLC38A2, NEAT1, FKBP5*, *ZBTB16*, *NR3C2*), and shortly thereafter module M2 (blue; *PLP1, CRYAB, CNP*; mature myelinating metabolic/proteostatic stress) rose when putative homeostatic motility/myelination modules (M5 & M7) began to decline (**Fig. 3e,f**).

Collectively, early pseudo-progression states were marked by an increased expression of genes associated with oligodendrocyte maturation and myelination (*MOG, MBP*), while exhibiting a decrease in oligo-neuronal synaptic exchange (*GRIN1*). In intermediate stages, however, there was an upregulation of protein degradation programs (*FKBP5, ZBTB16*) at the expense of those involved in myelination. In advanced stages, dysfunctional or compensatory myelination programs may potentially persist as tau pathology advances. Notably, the upregulation of *FKBP5*- and *CRYAB*-mediated chaperone and unfolded protein responses emerged as pivotal molecular events in the disease progression within oligodendrocytes.

### PSP neurons differentially regulate synaptic, proteostasis and glial interaction networks converging on ribosomal and survival dysfunction

To resolve transcriptional state transitions associated with neuronal tau cytopathology, we partitioned neuronal nuclei into Exc-ULN, Exc-DLN and Inh-Neu, and applied supervised pseudo-progression modeling using the semi-quantitative NFT load as the staging variable (**Fig. 4a,b**). While hdWGCNA networks were inferred on the transcriptomes of all neurons, we fitted the pseudo-progression models for each neuronal cell type separately. This approach enabled cross-neuron comparisons of expression networks while allowing some granularity for individual neuron cell types to be aligned with the pathology-anchored trajectory.

**Figure 4.**
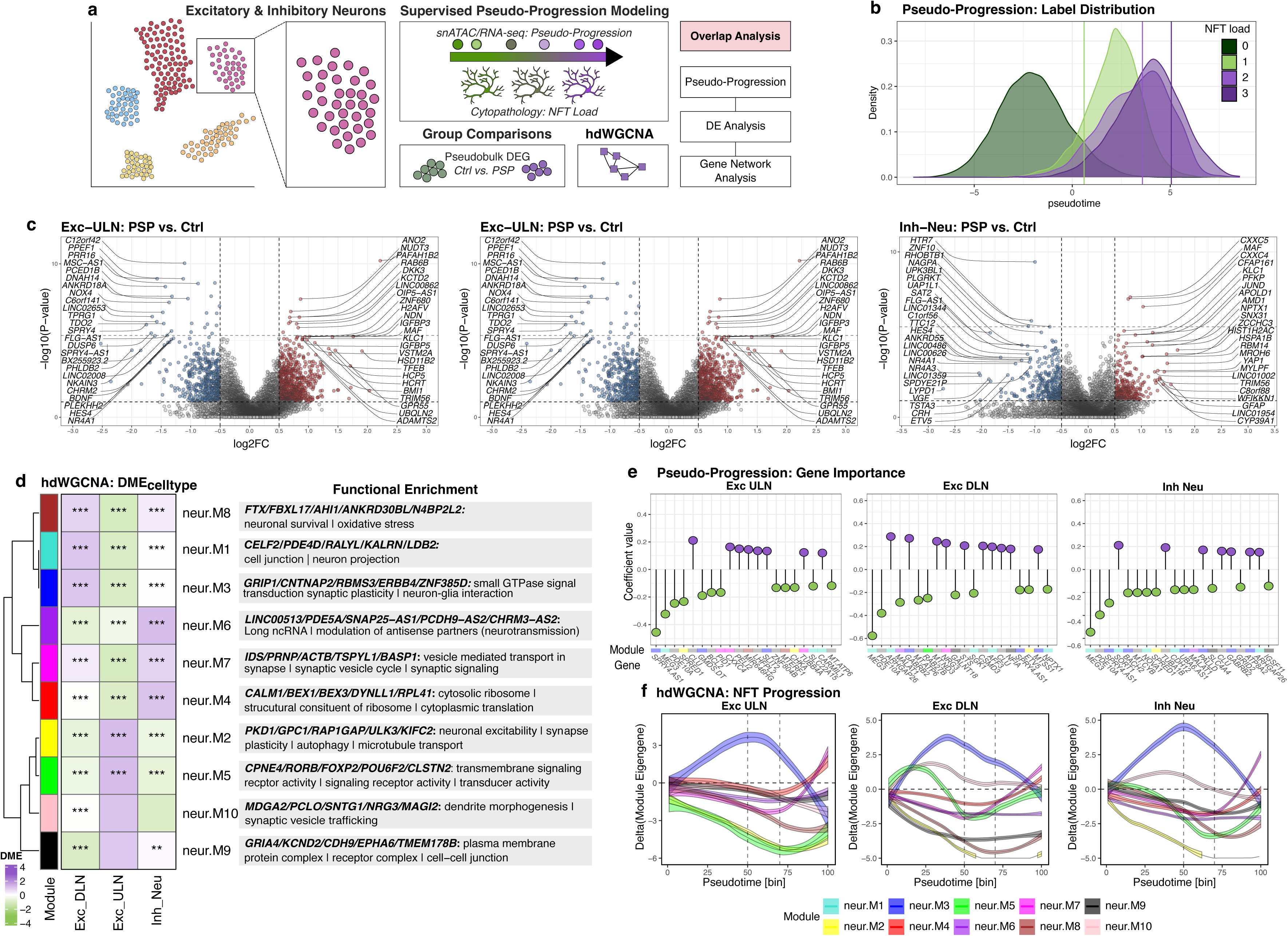
Neuronal transcriptional reprogramming along NFT pathology trajectories in PSP. **(a)** Analytical framework concept starting with the selection of Exc-ULN, Exc-DLN and Inh-Neu, and association testing of omic data with increasing NFT cytopathology burden, i.e. modelling of ‘pseudo-progression’. In parallel, group comparisons through pseudo-bulk DEG and cell type-resolved WGCNA analyses were conducted. **(b)** Density plots of NFT load labels over estimated pseudo-progression. **(c)** Volcano plot from pseudo-bulk DEG testing comparing PSP and control snRNAseq neurons (Exc-ULN, Exc-DLN and Inh-Neu) data. Top 25 up- and downregulated genes are labelled. **(d)** Heatmap displaying DME of PSP and control snRNAseq neuronal class data, stratified by neuronal cell type and 10 distinct hdWGCNA modules (neur.M1-neur.M10). Right-hand panels indicate GO enrichment of each module; top 5 hub genes and top functional enrichment terms are indicated. **(e)** Lollipop plots of neuronal gene expression pseudo-progression associations, separately for each cell type (Exc-ULN, Exc-DLN and Inh-Neu). Coefficient values of single genes are indicated, as provided from the final NFT model fit. Color tiles at the bottom indicate hdWGCNA module assignment. **(f)** Loess fit of hdWGCNA module eigengene values over pseudotime (i.e. pseudo-progression), separately for each cell type (Exc-ULN, Exc-DLN and Inh-Neu). Delta values are provided relative to the baseline. Color indicates neuronal hdWGCNA modules. <u>Abbreviations</u>: control (Ctrl), differentially expressed genes (DEG), differential module eigengene (DME), excitatory deep-layer neurons (Exc-DLN), excitatory upper-layer neurons (Exc-ULN), gene ontology (GO), high-dimensional weighted gene co-expression network analysis (hdWGCNA), inhibitory neurons (Inh-Neu), log2-fold-change (log2FC), neurofibrillary tangle (NFT).

Cell type-level DEG analysis of neuron populations showed partially distinct patterns (**Fig. 4c, Extended Data Fig. 8**). In Exc-ULN this highlighted significant upregulation of the Ca^2+^ channel *ANO2/TMEM16B*, the master regulator of autophagy *TFEB*, and the microtubule transport protein *RAB6B.* Conversely, genes related to reduced oxidative stress, neuronal survival and synaptic plasticity, such as *DUSP6* and *BDNF*, were downregulated. This pattern was partially mirrored in Exc-DLN with reduced expression of *BDNF* and the synaptic structural gene *CNTN6*, on the contrary there was increased expression of proteostasis-related genes (*HSP90AA1*, *HSP90AB1*, *NR1D1*). In Inh-Neu, DEG patterns were more distinct, featuring reduced synapse genes *VGF* and *HTR7,* as well as increased *CXXC5* – a negative feedback regulator of Wnt/β-catenin. Notably, PSP Inh-Neu also upregulated the canonical astrocytic cytoarchitecture protein *GFAP*; a neuronal isoform has been characterized before [42].

Pseudotime feature importance analysis identified genes associated directly with NFT cytopathology progression, including the cross-neuronal downregulation of long non-coding RNA *MEG3* and phosphodiesterase *PDE10A,* alongside shared up-regulation of the chaperone *CLU* (**Fig. 4e**). Several cell type-specific gene-pseudo-progression associations mainly centered on cytoskeletal remodeling (*CARMIL1, MTSS1*, *TUBB4A/TUBA1B*) and synaptic reorganization (*NPTX1*, *RAB27B, IGSF11, GABRB2*).

Gene network analysis using hdWGCNA identified ten pan-neuronal modules (neur.M1-neur.M10), exhibiting shared and unique transcriptional patterns across neuronal subtypes with increasing NFT burden. (**Fig. 4d-f, Extended Data Fig. 11**). While excitatory neuron subtypes broadly shared module dynamics, Exc-DLN and Inh-Neu exhibited even greater similarity. This suggests that adaptive mechanisms during NFT progression diverge from traditional neurochemical-based neuronal classification.

Module neur.M3 (blue; neuron-glia interaction) displayed biphasic dynamics across all neurons, rising to a peak at mid-stage pseudo-progression before a steady decline. In contrast, modules neur.M2 (yellow; neuronal excitability, autophagy, microtubule transport), neur.M8 (red; neuronal survival and oxidative stress), neur.M6 (purple; neurotransmission-regulatory ncRNAs), and neur.M9 (black; plasma membrane complexes) showed an early decline across all neurons. Modules neur.M8 and neur.M9 exhibited the most pronounced loss in Exc-DLN.

Module neur.M5 (green; transmembrane signaling) displayed marked cell type-specific divergence: it declined steadily in Exc-ULN, but in Exc-DLN had biphasic up- followed by downregulation. In Inh-Neu, neur.M5 decreased early. This reflects neuron subtype-dependent regulation of external signaling integration. Similarly, module neur.M10 (pink; dendrite morphogenesis and synaptic vesicle trafficking) demonstrated continuous slow decline in Exc-ULN, whereas both Exc-DLN and Inh-Neu displayed comparable biphasic up-down trajectories.

Finally, modules neur.M4 (red; cytosolic ribosome and cytoplasmic translation) and neur.M7 (magenta; synaptic vesicle cycle) increased exclusively during advanced-stage pseudo-progression across all neuronal cell types. This indicates convergent activation of vesicular trafficking and ribosomal stress-response pathways during advanced NFT cytopathology (**Fig. 4f**).

### Cross-modal concordance between brain transcriptomics and CSF proteomics

To assess the external validity of the cell type-specific transcriptional alterations, we intersected snRNAseq DEGs from our PSP cohort with differentially abundant proteins from an independent PSP CSF proteomics dataset [37]. Pearson correlation between brain transcript and CSF protein fold-changes was cell type-dependent and overall modest (**Fig. 5a**), consistent with attenuation across the brain-CSF barrier and heterogeneous protein secretion profiles, reaching statistical significance only in Inh-Neu (R = 0.12, p = 2.2×10⁻) and OPC (R = 0.058, p = 0.041), with a weak, trending positive correlation in Astro (R = 0.05, p = 0.071). Notably, global fold-change correlation and concordant gene-protein pair fractions were partially dissociated: Exc-DLN, Inh-Neu, and Oligo exhibited the highest proportion of significant-concordant pairs (>60%; **Fig. 5b**) despite a non-significant aggregate correlation in Exc-DLN, suggesting selective convergence of a biologically coherent target subset rather than uniform transcriptome-proteome coupling. The most robustly concordant targets across cell types included markers of glial activation (C*HI3L1, GFAP*), neuroaxonal damage (*NEFM*), and synaptic function (*VGF, NPTX2, LYPD1;* **Fig. 5c**), nominating these as priority biomarker candidates reflecting PSP-associated pathophysiology in both brain tissue and CSF.

**Figure 5.**
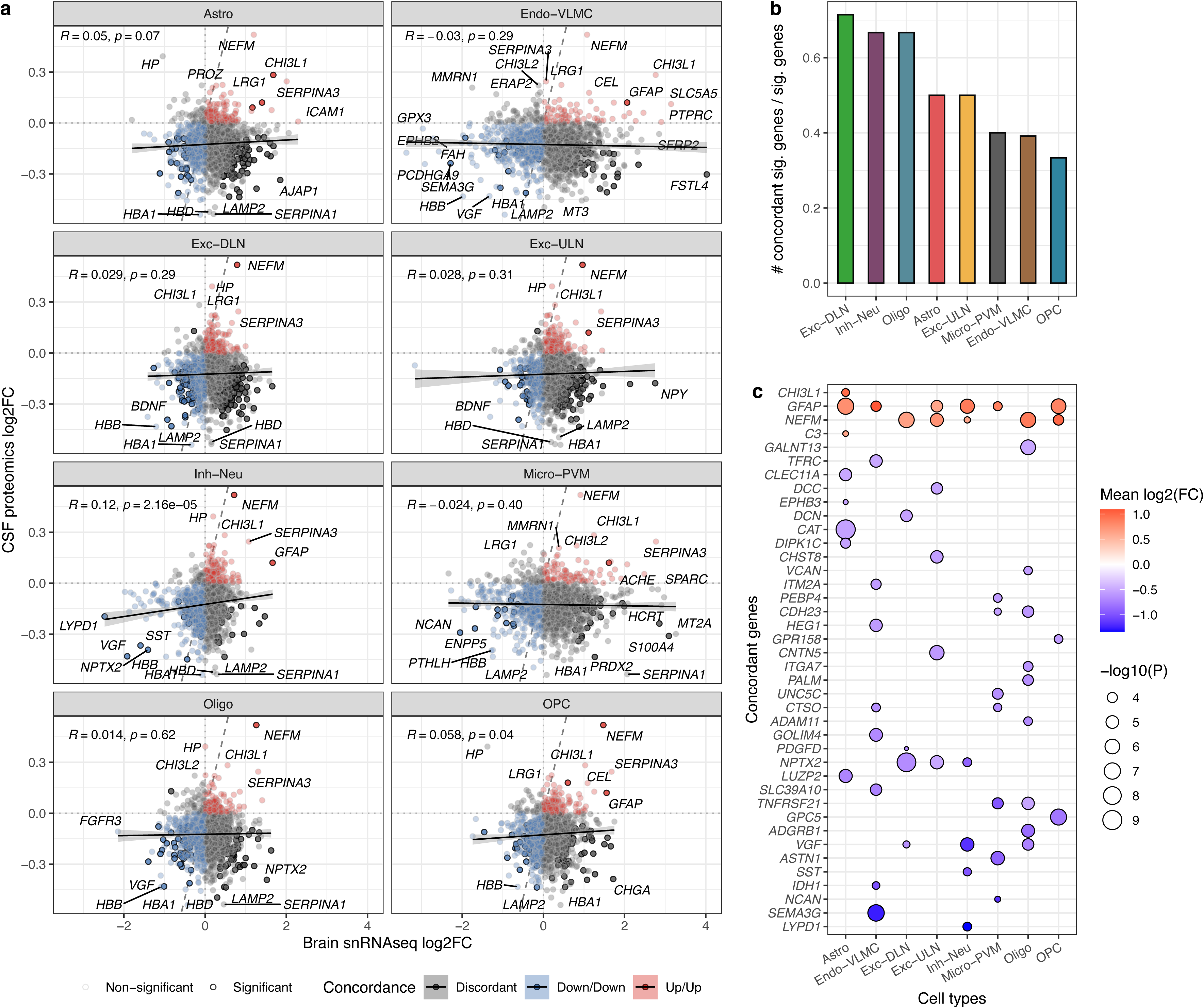
Cross-modal concordance between brain transcriptomic and CSF proteomic changes in PSP. **(a)** Scatter plots of brain snRNAseq log2 fold-change (x-axis, PSP vs. control) vs. CSF proteomics log2FC (y-axis, PSP vs. HC) for all genes detected in both datasets, faceted by cell type. Each point represents one gene; color encodes directional concordance (red: up/up; blue: down/down; grey: discordant); black encircled points/candidates reached nominal significance in snRNAseq and CSF proteomics (p < 0.05). The dashed diagonal line represents identity (slope = 1). Pearson R and p-value are shown per facet. **(b)** Bar plot showing the fraction of statistically significant (snRNAseq and CSF proteomics p < 0.05) DEG-cell type pairs with concordant direction of regulation between brain transcriptomics and CSF proteomics, across the eight analysed cell types. **(c)** Dot plot of concordant DEG-cell type pairs meeting both significance and effect-size thresholds (snRNAseq and CSF proteomics p < 0.05; |mean log2FC| > 0.5). Point fill encodes mean log2FC across both modalities (red: upregulated; blue: downregulated in PSP); point size encodes −log10(snRNAseq p-value). <u>Abbreviations</u>: cerebrospinal fluid (CSF), healthy control (HC), log2-fold-change (log₂FC), significant (sig.).

## Discussion

In this work, we present the first integrated single-nucleus transcriptomic-epigenetic dataset of PSP frontal cortex, comprising over 144,000 nuclei of eight PSP and eight control cases and integrating molecular profiles with semi-quantitative tau cytopathology. Our multimodal approach revealed cell type-specific vulnerabilities, with PSP cases exhibiting preferential loss of deep-layer neuron and glial subclusters relative to inhibitory neurons. Although only nominally significant and exploratory, GWAS variant enrichment analysis points toward a glial-centered genetic susceptibility, with PSP risk variants most enriched in astrocytes and oligodendrocytes. Cross-cell-type comparisons of gene expression demonstrated that excitatory neurons harbor the greatest burden of differentially expressed genes, characterized by upregulated proteostasis and downregulated synaptic-neuronal integrity genes. The pseudo- progression modeling provides a tau histopathology-centric approach to understanding the distinct temporal trajectories of PSP pathophysiology, suggesting potential stage-specific therapeutic windows.

Our finding of glial-centered genetic susceptibility builds upon our previous GWAS integration analysis, which demonstrated enrichment of PSP risk variants in astrocytes [13], and now extends this finding to oligodendrocytes. The largest PSP GWAS comprising 2’779 cases confirmed significant loci at *MAPT, MOBP, STX6, RUNX2, SLCO1A2* and *C4A* [43], with others identifying additional susceptibility loci including *SP1* [11], *KIF13A,* and *DUSP10* [44]. The only CBD GWAS identified overlapping loci at *MOBP*, *KIF13B* and *DUSP4* among few others [10], reinforcing the shared genetic architecture between these 4R tauopathies. Several dysregulated candidates in our analysis correspond to established risk genes or are paralogs thereof: *MOBP* (biphasic dysregulation in oligodendrocytes), SP1 (TF upregulated in Micro-PVM), *KIF2C* (neuronal microtubule dynamics), *DUSP6/DUSP3* (dysregulated in excitatory neurons and astrocytes). Additionally, the AD risk gene and chaperone *CLU* was upregulated in intermediate NFT burden in neurons in our dataset, reflecting a pan-neuronal proteostasis response. Beyond SP1 as a deregulated TF in PSP microglia, our analysis did not confirm microglial risk variant enrichment, which contrasts with the genetic enrichment observed in AD, and suggests distinct immunological mechanisms in PSP compared to AD. Together with the high DEG burden in excitatory neurons (followed by astrocytes and oligodendrocytes), this implies that although genetic risk may be determined by these glial populations, transcriptomic dysregulation and downstream neurodegeneration apparently act through multiple lineages in PSP.

The pseudo-progression modeling approach, anchored in cell type-specific tau pathology, offers hypothetical trajectories of molecular reprogramming (as detailed in **Fig. 6**). In oligodendrocytes, co-expression networks shifted from homeostatic myelination to stress signaling as the CB burden increased (**Fig. 6b**). Significant and potentially druggable targets may involve promoting early sustained myelination and oligodendrocytic-neuronal signaling through *MBP/MOBP* and gliotransmission via ephrin receptors, respectively. The strongest positive association with disease progression was observed for *FKBP5* expression, which encodes FK506-binding protein 51 – an immunophilin and co-factor of the 90 kDa heat-shock protein (Hsp90). FKBP5 forms a mature chaperone complex with Hsp90 that prevents effective tau degradation and promotes tau oligomerization [45,46]. Homologue-specific FKBP5 inhibitors, such as SAFit2 [47], have demonstrated neuroprotective effects in PD mouse models, reducing ubiquitin-positive inclusions, neurodegeneration and motor dysfunction [48]. Our discovery that *FKBP5* expression in oligodendrocytes is associated with disease progression broadens this mechanism to glial tau pathology and indicates *FKBP5* as a (non-disease-specific) marker of oligodendrocytic CB burden in PSP. Although the specific role of *FKBP5* in oligodendrocytes remains largely unexplored, elevated FKBP5 levels may hinder proteasomal tau clearance, thereby diminishing myelination capabilities and axonal support. Consequently, isoform-selective inhibition of FKBP5 could represent a viable therapeutic target for PSP as well. Additionally, UCHL1, a deubiquitinating enzyme previously found in glial cytoplasmic inclusions in multiple system atrophy (MSA) [49], was significantly upregulated in Oligo at high CB burden in our dataset, potentially indicating shared mechanisms of inclusion formation across α- synucleinopathies and 4R tauopathies.

**Figure 6.**
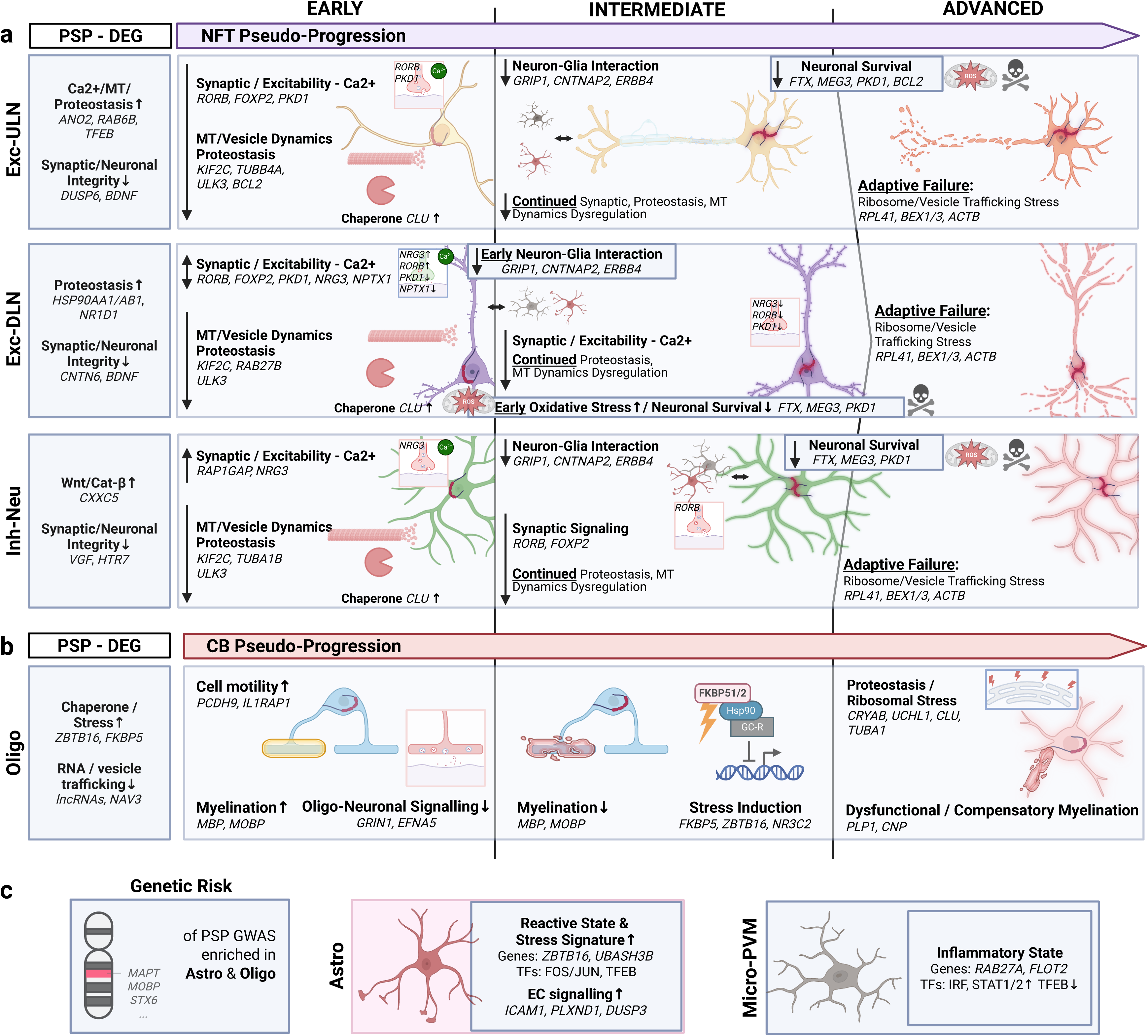
Conceptual framework of cell type-specific pathological changes in PSP. Schematic representation of differential expression and molecular events associated with tau cytopathology pseudo- progression across neuronal and glial lineages. **(a)** In Inh-Neu and excitatory (Exc-DLN, Exc-ULN) neurons, early stages are defined by dysregulation of synaptic excitability, Ca²⁺ signaling, proteostasis, and MT/vesicle dynamics. Intermediate progression involves divergent neuron-glia interactions, ROS, and survival pathways, which ultimately converge in an advanced state of adaptive failure characterized by ribosomal stress, impaired vesicle trafficking, and diminished survival signaling. (**b)**. Oligo pathology transitions from early functional myelination and oligo-neuronal signaling to a *FKBP5*-driven stress signature and defective myelination, ultimately reaching a late-stage state of proteostatic and ribosomal stress alongside maladaptive or sustained compensatory myelination. **c**. Genetic risk is primarily enriched in Astro and Oligo. Astro and Micro-PVM lineages demonstrate distinct, stage-dependent reactive and inflammatory states. Created with BioRender. <u>Abbreviations</u>: astrocytes (Astro), calcium (Ca²⁺), coiled body (CB), differentially expressed genes (DEG), excitatory deep-layer neurons (Exc-DLN), excitatory upper-layer neurons (Exc-ULN), extracellular signaling (EC), genome-wide association study (GWAS), inhibitory neurons (Inh-Neu), microglia and perivascular macrophages (Micro-PVM), microtubule (MT), neurofibrillary tangle (NFT), oligodendrocytes (Oligo), reactive oxygen species (ROS), transcription factors (TFs).

Across neuronal cell types, co-expression modules revealed differential patterns, involving early dysregulation of neuronal, structural and functional integrity pathways, differently timed neuron-glia interactions and oxidative stress signals, followed by late-stage convergence on adaptive failure with reduced neuronal survival cues (**Fig. 6a**). Interestingly, the neuron-glia interaction and oxidative stress modules were tended to be altered earlier in Exc-DLN, implying higher vulnerability or mirroring the preferential deep-layer involvement of PSP-type tau pathology observed in early neuropathological studies [50]. Exc-ULN and Exc-DLN had the highest DEG burden among all cell types and were characterized by coordinated upregulation of proteostasis machinery (*HSP90, TFEB*) and downregulation of synaptic integrity genes (*CNTN6*, *DUSP6*, *BDNF*). Critical molecular targets implicate restoring depleted BDNF levels, and neuron-subtype-aware replenishment of synaptic-hyperexcitability centered around *RORB, NRG3 and NPTX1*.

RORB selectively marks vulnerable excitatory neurons in the entorhinal cortex and deep cortical layers, where these neurons preferentially accumulate NFTs and undergo early degeneration in AD [7]. This vulnerability might extend to synaptic damage in PSP, through RORB-mediated transcriptional regulation, as well as NRG3 and NPTX1 networks. Downregulating *NRG3* in excitatory neurons, which promotes excitatory synapse formation in interaction with ErbB4+ Pvalb interneurons [51], could weaken inhibitory circuit strength and amplify the hyperexcitability observed in the progression analysis. Additionally, reduced expression of *NPTX2*, which restrains complement-1q-mediated synaptic pruning, would combine with tau-driven effects to diminish synaptotrophic support. Restoring *NRG3* and *NPTX2* in vulnerable neuronal populations offers another potential therapeutic option to stabilize synaptic circuits.

Similarly to *FKBP5* in oligodendrocytes, the expression of *clusterin* (*CLU*) started to increase across neurons during intermediate progression stage, indicating a potential tau-amplifying or failed compensatory upregulation. CLU’s function is context-dependent: its chaperone activity appears to follow a CLU-to-tau ratio-dependent manner, wherein it can delay tau fibril formation at higher ratios while paradoxically enhancing oligomeric tau seeding capacity at lower ratios [52]. These properties may ultimately contribute to intercellular tau propagation through endo-lysosomal escape and extracellular release mechanisms. While CLU function restoration is being investigated in AD patients with *CLU* loss-of-function mutations [53], its role in 4R tauopathies – whether detrimental or protective – requires further investigation.

Preceding *CLU* upregulation, the concomitant dysregulation of proteostasis and Ca²⁺ homeostasis factors (*ULK3, HSP90AA1/AB1, TFEB, ANO2*) in neurons, alongside microtubule dynamics regulators (*KIF2C, TUBA1B, TUBB4A*) and vesicle trafficking genes (*RAB27B*), may reflect compensatory attempts to restore axonal transport disrupted by tau-microtubule dissociation. However, progressive failure of these proteostasis pathways is evidenced by ribosomal stress markers (*RPL41, BEX1/3*) emerging at intermediate to advanced stages, suggesting maladaptive responses culminating in neurodegeneration.

Ca²⁺ dysregulation could represent a convergent pathogenic mechanism across neuronal cell types and tauopathies. In genetic Drosophila tauopathy models, pathogenic tau depletes nuclear Ca²⁺ and CREB to drive neuronal death [54]. Conversely, activation of BK (big potassium) channels elevates nuclear Ca²⁺ levels and suppresses tau neurotoxicity. Similarly, Tau oligomer-mediated excitotoxicity through glutamatergic receptors can be reduced by the targeted knockdown of a potassium channel subunit, which also decreases tau oligomer formation [55]. These findings nominate Ca²⁺ dynamics as potential therapeutic targets for excitatory neuron protection in PSP.

A major common theme across both cell classes, neurons and oligodendrocytes, was the downregulation of *MEG3* (a long non-coding RNA gene) association with either NFT or CB progression. Notably, the role of *MEG3* also appears to be context-dependent: In PD models, *MEG3* downregulation leads to apoptosis through reduced *LRRK2* expression [56]. Conversely, post-ischemic upregulation promotes neuronal apoptosis [57], while in chronic neurodegenerative conditions such as AD, it may exert neuroprotective effects by modulating the PI3K/Akt pathway [58]. This variable functional profile indicates that *MEG3* exerts context-dependent effects that depend on both the neurodegenerative condition and the stage of disease progression.

While excitatory neurons had the greatest DEG burden, inhibitory neurons showed strongest TF deregulation extents and partly different alterations along the tau pseudo-progression trajectory. A key distinction was *CXXC5* upregulation in DEG analysis, which resembles the situation in AD-type amyloidosis models [59], where it suppresses Wnt/β- catenin signaling activation and leads to neuronal apoptosis. Inhibitory neurons also exhibited lower expression of distinct synaptic and neuronal integrity genes (*VGF, HTR7*), suggesting impaired neurotransmitter signaling; notably, serotonin receptor *HTR7* reduction has been associated with cognitive-neuropsychiatric deficits in AD [60]. Curiously, the increased expression of *GFAP* in inhibitory neurons has been described before, where it might reflect the overexpression of a neuronal isoform [42]. Despite the notable differences from excitatory neurons, the dysregulation of genes involved in Ca²⁺ signaling and neuronal hyperexcitability, microtubule and vesicle dynamics, as well as proteostasis largely resembles that of excitatory upper-layer neurons. Despite transcriptional dysregulation present, the relatively moderate extent in inhibitory neurons point to either cell-type-specific resilience in the studied 4R tauopathy or pathomechanisms that remain undetected in this dataset.

Integrating cortical transcriptomics with an independent PSP CSF proteomics dataset [37] revealed that only a select subset of transcriptional changes is reflected at the CSF protein level. The globally modest and cell type-dependent concordance is consistent with an expected attenuation across the brain and CSF compartments, and likely reflects regional heterogeneity of protein origin, differential release into the extracellular space, and dilution by proteomic sources other than the frontal cortex. Significant aggregate fold-change correlations were restricted to inhibitory neurons, OPC and weakly in astrocytes. Yet several neuronal and glial cell types showed >60% individually concordant gene-protein pairs, indicating that the transcriptome-proteome relationship is driven by a high-signal target subset rather than a uniform shift. The large discordant fraction likely reflects compartment specificity or intracellularly enriched transcripts inaccessible to CSF proteomics. However, the convergent protein targets recapitulate pathway alterations implicated by our study with markers of glial reactivity (*CHI3L1, GFAP*), axonal injury (*NEFM*) and synaptic function (*VGF, NPTX2, LYPD1*). These markers are biologically traceable to CSF, nominating key targets for prospective fluid biomarker validation.

In summary, this study provides a conceptual framework in which genetic predisposition primarily operates through glial dysfunction, while epigenetic-transcriptomic dysregulation occurs across multiple cell lineages, and along distinct temporal trajectories. However, it is important to emphasize, that these pseudo-progression alterations likely reflect complex multicellular dynamics rather than deterministic cell-autonomous trajectories. While the supervised pseudotime analysis orders cells along histopathology-correlated transcriptional axes, this ordering may arise from at least two non-exclusive mechanisms: (i) cell state shifts within persistent cell populations, or (ii) selective depletion of specific subclusters coupled with expansion or preservation of others. Additionally, tau pathology may exert non-cell- autonomous effects within local microenvironments, whereby a high local soluble tau burden likely influences neighboring cells regardless of their intrinsic insoluble tau load. Consequently, we interpret pseudo-progression as a histological community-level trajectory – i.e., a population-level readout of coordinated but heterogeneous single-cell fates – wherein some cells undergo transcriptional state shifts while others would degenerate and are lost from the analyzed population. This multicellular framework, consistent with recent studies in AD [17], suggests that PSP pathogenesis unfolds through coordinated shifts across interacting cell populations.

### Limitations and Future Directions

Several limitations warrant consideration. First, our cross-sectional design semi-quantitative tau staging in a comparably small cohort provides inferred, not actual longitudinally sampled, disease trajectories. Prospective biomarker studies of larger cohorts correlating *antemortem* CSF/imaging measures with *postmortem* molecular profiles would strengthen causal inference and improve generalizability. Second, snATACseq and snRNAseq represent *nuclear* transcriptomes and chromatin states, potentially missing cytoplasmic RNA populations and post- transcriptional regulatory mechanisms critical to tau pathobiology. Furthermore, tissue artifacts due to agony cannot be excluded, though where possible we included the postmortem interval as covariate. Third, the focus on frontal cortex, though clinically relevant, limits generalizability to subcortical regions (e.g., basal ganglia, subthalamic nucleus, substantia nigra) where PSP pathology is equally or even more prominent. Fourth, while our cohort matching minimized confounding neuropathologies, subtle AD co-pathology in some cases may influence interpretations; single-pathology PSP cohorts are exceedingly rare but would provide greater specificity. Future directions should integrate spatial multi-omics technologies that preserve tissue architecture while achieving single-cell resolution, enabling precise mapping of tau pathology to transcriptional states within cytoarchitectonic layers and anatomical subregions. Integration of proteomic and metabolomic data would bridge the gap between gene expression and functional cellular phenotypes. Finally, translating candidate therapeutic targets identified here into *in vivo* models – particularly those recapitulating oligodendrocytic and neuronal tau pathology – will be essential for preclinical validation and mechanistic studies.

## Acknowledgements

We thank the donors and their families for making this research possible, the associates of the Neurobiobank Munich for structural support, and Dr. N. Buresch for technical support.

## References

1. Höglinger GU, Respondek G, Stamelou M, Kurz C, Josephs KA, Lang AE, et al. Clinical diagnosis of progressive supranuclear palsy: The movement disorder society criteria. Movement Disorders [Internet]. John Wiley & Sons, Ltd; 2017 [cited 2019 Jun 7];32:853–64. 10.1002/mds.26987

2. Narasimhan S, Guo JL, Changolkar L, Stieber A, McBride JD, Silva L V., et al. Pathological Tau Strains from Human Brains Recapitulate the Diversity of Tauopathies in Nontransgenic Mouse Brain. The Journal of Neuroscience [Internet]. 2017;37:11406–23. 10.1523/JNEUROSCI.1230-17.2017

3. Shi Y, Zhang W, Yang Y, Murzin AG, Falcon B, Kotecha A, et al. Structure-based classification of tauopathies. Nature 2021 598:7880 [Internet]. Nature Publishing Group; 2021 [cited 2024 Feb 2];598:359–63. 10.1038/s41586-021-03911-7

4. Kovacs GG. Astroglia and Tau: New Perspectives. Front. Aging Neurosci. Frontiers Media S.A.; 2020. 10.3389/fnagi.2020.00096

5. Ferrer I, López-González I, Carmona M, Arregui L, Dalfó E, Torrejón-Escribano B, et al. Glial and Neuronal Tau Pathology in Tauopathies. J Neuropathol Exp Neurol [Internet]. Narnia; 2014 [cited 2019 Aug 29];73:81–97. 10.1097/NEN.0000000000000030

6. Mathys H, Boix CA, Akay LA, Xia Z, Davila-Velderrain J, Ng AP, et al. Single-cell multiregion dissection of Alzheimer’s disease. Nature. 2024; 10.1038/s41586-024-07606-7

7. Leng K, Li E, Eser R, Piergies A, Sit R, Tan M, et al. Molecular characterization of selectively vulnerable neurons in Alzheimer’s disease. Nat Neurosci [Internet]. Nature Publishing Group; 2021 [cited 2021 Jan 21];1–12. 10.1038/s41593-020-00764-7

8. Briel N, Pratsch K, Roeber S, Arzberger T, Herms J. Contribution of the astrocytic tau pathology to synapse loss in progressive supranuclear palsy and corticobasal degeneration. Brain Pathology [Internet]. Wiley; 2020 [cited 2020 Dec 23]; 10.1111/bpa.12914

9. Richetin K, Steullet P, Pachoud M, Perbet R, Parietti E, Maheswaran M, et al. Tau accumulation in astrocytes of the dentate gyrus induces neuronal dysfunction and memory deficits in Alzheimer’s disease. Nature Neuroscience 2020 23:12 [Internet]. Nature Publishing Group; 2020 [cited 2022 Dec 23];23:1567–79. 10.1038/s41593-020-00728-x

10. Kouri N, Ross OA, Dombroski B, Younkin CS, Serie DJ, Soto-Ortolaza A, et al. Genome-wide association study of corticobasal degeneration identifies risk variants shared with progressive supranuclear palsy. Nat Commun. 2015;6:1–7. 10.1038/ncomms8247

11. Chen JA, Chen Z, Won H, Huang AY, Lowe JK, Wojta K, et al. Joint genome-wide association study of progressive supranuclear palsy identifies novel susceptibility loci and genetic correlation to neurodegenerative diseases. Mol Neurodegener. BioMed Central Ltd.; 2018;13. 10.1186/s13024-018-0270-8

12. Höglinger GU, Melhem NM, Dickson DW, Sleiman PMA, Wang L-S, Klei L, et al. Identification of common variants influencing risk of the tauopathy progressive supranuclear palsy. Nat Genet [Internet]. Nature Publishing Group; 2011 [cited 2019 Oct 7];43:699–705. 10.1038/ng.859

13. Briel N, Ruf VC, Pratsch K, Roeber S, Widmann J, Mielke J, et al. Single-nucleus chromatin accessibility profiling highlights distinct astrocyte signatures in progressive supranuclear palsy and corticobasal degeneration. Acta Neuropathol [Internet]. Acta Neuropathol; 2022 [cited 2023 Jun 3];144:615–35. 10.1007/S00401-022-02483-8

14. Mukherjee S, Klaus C, Pricop-Jeckstadt M, Miller JA, Struebing FL. A microglial signature directing human aging and neurodegeneration-related gene networks. Front Neurosci [Internet]. Frontiers Media S.A.; 2019 [cited 2021 May 20];13:2. 10.3389/fnins.2019.00002

15. Whitney K, Song WM, Sharma A, Dangoor DK, Farrell K, Krassner MM, et al. Single-cell transcriptomic and neuropathologic analysis reveals dysregulation of the integrated stress response in progressive supranuclear palsy. Acta Neuropathol. Springer Science and Business Media Deutschland GmbH; 2024;148. 10.1007/s00401-024-02823-w

16. Rexach JE, Cheng Y, Chen L, Polioudakis D, Lin LC, Mitri V, et al. Cross-disorder and disease-specific pathways in dementia revealed by single-cell genomics. Cell. Elsevier B.V.; 2024;187:5753–5774.e28. 10.1016/j.cell.2024.08.019

17. Green GS, Fujita M, Yang HS, Taga M, Cain A, McCabe C, et al. Cellular communities reveal trajectories of brain ageing and Alzheimer’s disease. Nature. Nature Research; 2024;633:634–45. 10.1038/s41586-024-07871-6

18. Stamelou M, Respondek G, Giagkou N, Whitwell JL, Kovacs GG, Höglinger GU. Evolving concepts in progressive supranuclear palsy and other 4-repeat tauopathies. Nat Rev Neurol. Nature Research; 2021;17:601–20. 10.1038/S41582-021-00541-5

19. Valletta M, Briel N, Yuksekel I, Barboure M, Coward A, De Houwer JFH, et al. Fluid biomarkers for neurodegenerative diseases: a comprehensive update. Alzheimers Res Ther. 2025; 10.1186/s13195-025-01919-z

20. Hauw JJ, Daniel SE, Dickson D, Horoupian DS, Jellinger K, Lantos PL, et al. Preliminary NINDS Neuropathologic Criteria for Steele-Richardson-Olszewski Syndrome (Progressive Supranuclear Palsy). Neurology. Wolters Kluwer Health, Inc. on behalf of the American Academy of Neurology; 1994. p. 2015–9. 10.1212/wnl.44.11.2015

21. Litvan I, Hauw JJ, Bartko JJ, Lantos PL, Daniel SE, Horoupian DS, et al. Validity and Reliability of the Preliminary NINDS Neuropathologic Criteria for Progressive Supranuclear Palsy and Related Disorders. J Neuropathol Exp Neurol [Internet]. Lippincott Williams and Wilkins; 1996 [cited 2020 Mar 3];55:97–105. 10.1097/00005072-199601000-00010

22. Braak H, Del Tredici K, Rüb U, De Vos RAI, Jansen Steur ENH, Braak E. Staging of brain pathology related to sporadic Parkinson’s disease. Neurobiol Aging. 2003;24:197–211. 10.1016/S0197-4580(02)00065-9

23. Hao Y, Stuart T, Kowalski MH, Choudhary S, Hoffman P, Hartman A, et al. Dictionary learning for integrative, multimodal and scalable single-cell analysis. Nat Biotechnol. Nature Research; 2024;42:293–304. 10.1038/s41587-023-01767-y

24. Stuart T, Srivastava A, Madad S, Lareau CA, Satija R. Single-cell chromatin state analysis with Signac. Nat Methods. Nature Research; 2021;18:1333–41. 10.1038/s41592-021-01282-5

25. Korsunsky I, Millard N, Fan J, Slowikowski K, Zhang F, Wei K, et al. Fast, sensitive and accurate integration of single-cell data with Harmony. Nature Methods 2019 16:12 [Internet]. Nature Publishing Group; 2019 [cited 2022 Aug 19];16:1289–96. 10.1038/s41592-019-0619-0

26. Castro-Mondragon JA, Riudavets-Puig R, Rauluseviciute I, Berhanu Lemma R, Turchi L, Blanc-Mathieu R, et al. JASPAR 2022: The 9th release of the open-access database of transcription factor binding profiles. Nucleic Acids Res. Oxford University Press; 2022;50:D165–73. 10.1093/nar/gkab1113

27. Schep AN, Wu B, Buenrostro JD, Greenleaf WJ. ChromVAR: Inferring transcription-factor-associated accessibility from single-cell epigenomic data. Nat Methods [Internet]. Nature Publishing Group; 2017 [cited 2021 Jan 19];14:975–8. 10.1038/nmeth.4401

28. caleblareau/gchromVAR: Cell type specific enrichments using finemapped variants and quantitative epigenetic data [Internet]. [cited 2020 Apr 3]. https://github.com/caleblareau/gchromVAR. Accessed 3 Apr 2020

29. Hao Y, Hao S, Andersen-Nissen E, Mauck WM, Zheng S, Butler A, et al. Integrated analysis of multimodal single-cell data. Cell. Elsevier B.V.; 2021;184:3573–3587.e29. 10.1016/j.cell.2021.04.048

30. Grabski IN, Street K, Irizarry RA. Significance analysis for clustering with single-cell RNA-sequencing data. Nat Methods. Nature Research; 2023;20:1196–202. 10.1038/s41592-023-01933-9

31. Mangiolaa S, Schulzea AJR, Trussarta M, Valdesa EZ, Maa M, Gaoc Z, et al. Sccomp: Robust differential composition and variability analysis for single-cell data. Proc Natl Acad Sci U S A. National Academy of Sciences; 2023;120. 10.1073/pnas.2203828120

32. Macnair W, Gupta R, Claassen M. Psupertime: Supervised pseudotime analysis for time-series single-cell RNA- seq data. Bioinformatics. Oxford University Press; 2022;38:I290–8. 10.1093/bioinformatics/btac227

33. Love MI, Huber W, Anders S. Moderated estimation of fold change and dispersion for RNA-seq data with DESeq2. Genome Biol. BioMed Central Ltd.; 2014;15. 10.1186/s13059-014-0550-8

34. Morabito S, Reese F, Rahimzadeh N, Miyoshi E, Swarup V. hdWGCNA identifies co-expression networks in high- dimensional transcriptomics data. Cell Reports Methods. Cell Press; 2023;3. 10.1016/j.crmeth.2023.100498

35. Gu Z, Eils R, Schlesner M. Complex heatmaps reveal patterns and correlations in multidimensional genomic data. Bioinformatics. Oxford University Press; 2016;32:2847–9. 10.1093/bioinformatics/btw313

36. Wu T, Hu E, Xu S, Chen M, Guo P, Dai Z, et al. clusterProfiler 4.0: A universal enrichment tool for interpreting omics data. Innovation. Cell Press; 2021;2. 10.1016/j.xinn.2021.100141

37. Jang Y, Oh S, Hall AJ, Zhang Z, Tropea TF, Chen-Plotkin A, et al. Biomarker discovery in progressive supranuclear palsy from human cerebrospinal fluid. Clin Proteomics. BioMed Central Ltd; 2024;21. 10.1186/s12014-024-09507-3

38. Ritchie ME, Phipson B, Wu D, Hu Y, Law CW, Shi W, et al. Limma powers differential expression analyses for RNA-sequencing and microarray studies. Nucleic Acids Res. Oxford University Press; 2015;43:e47. 10.1093/nar/gkv007

39. Cerezo M, Sollis E, Ji Y, Lewis E, Abid A, Bircan KO, et al. The NHGRI-EBI GWAS Catalog: Standards for reusability, sustainability and diversity. Nucleic Acids Res. Oxford University Press; 2025;53:D998–1005. 10.1093/nar/gkae1070

40. Du S, Zheng H. Role of FoxO transcription factors in aging and age-related metabolic and neurodegenerative diseases. Cell Biosci. 2021;11:188. 10.1186/s13578-021-00700-7

41. Galuh S, Faught E, Klaassen I, Koorneef LL, Brinks J, van Dijk EHC, et al. The glucocorticoid receptor is affected by its target ZBTB16 in a dissociated manner. Journal of Endocrinology. BioScientifica Ltd.; 2025;266. 10.1530/JOE-24-0283

42. Hol EM, Roelofs RF, Moraal E, Sonnemans MAF, Sluijs JA, Proper EA, et al. Neuronal expression of GFAP in patients with Alzheimer pathology and identification of novel GFAP splice forms. Mol Psychiatry. 2003;8:786–96. 10.1038/sj.mp.4001379

43. Farrell K, Humphrey J, Chang T, Zhao Y, Leung YY, Kuksa PP, et al. Genetic, transcriptomic, histological, and biochemical analysis of progressive supranuclear palsy implicates glial activation and novel risk genes. Nature Communications . Nature Research; 2024;15. 10.1038/s41467-024-52025-x

44. Wang H, Chang TS, Dombroski BA, Cheng PL, Patil V, Valiente-Banuet L, et al. Whole-genome sequencing analysis reveals new susceptibility loci and structural variants associated with progressive supranuclear palsy. Molecular Neurodegeneration . BioMed Central Ltd; 2024;19. 10.1186/s13024-024-00747-3

45. Blair LJ, Nordhues BA, Hill SE, Scaglione KM, O’Leary JC, Fontaine SN, et al. Accelerated neurodegeneration through chaperone-mediated oligomerization of tau. Journal of Clinical Investigation. 2013;123:4158–69. 10.1172/JCI69003

46. Jinwal UK, Koren J, Borysov SI, Schmid AB, Abisambra JF, Blair LJ, et al. The Hsp90 cochaperone, FKBP51, increases tau stability and polymerizes microtubules. Journal of Neuroscience. 2010;30:591–9. 10.1523/JNEUROSCI.4815-09.2010

47. Gaali S, Kirschner A, Cuboni S, Hartmann J, Kozany C, Balsevich G, et al. Selective inhibitors of the FK506- binding protein 51 by induced fit. Nat Chem Biol. Nature Publishing Group; 2015;11:33–7. 10.1038/nchembio.1699

48. Garcia-Gomara M, Legarra-Marcos N, Serena M, Rojas-de-Miguel E, Espelosin M, Marcilla I, et al. FKBP51 inhibition ameliorates neurodegeneration and motor dysfunction in the neuromelanin-SNCA mouse model of Parkinson’s disease. Molecular Therapy. Cell Press; 2025;33:895–916. 10.1016/j.ymthe.2025.01.049

49. Pukaß K, Richter-Landsberg C. Inhibition of UCH-L1 in oligodendroglial cells results in microtubule stabilization and prevents α-synuclein aggregate formation by activating the autophagic pathway: implications for multiple system atrophy. Front Cell Neurosci. 2015;9:163. 10.3389/fncel.2015.00163

50. Verny M, Duyckaerts C, Agid Y, Hauw JJ. The significance of cortical pathology in progressive supranuclear palsy. Clinico-pathological data in 10 cases. Brain. Oxford University Press; 1996;119:1123–36. 10.1093/brain/119.4.1123

51. Müller T, Braud S, Jüttner R, Voigt BC, Paulick K, Sheean ME, et al. Neuregulin 3 promotes excitatory synapse formation on hippocampal interneurons. EMBO J. Springer Science and Business Media LLC; 2018;37. 10.15252/embj.201798858

52. Yuste-Checa P, Trinkaus VA, Riera-Tur I, Imamoglu R, Schaller TF, Wang H, et al. The extracellular chaperone Clusterin enhances Tau aggregate seeding in a cellular model. Nat Commun. Nature Research; 2021;12. 10.1038/s41467-021-25060-1

53. Cohn W, Campagna J, Wi D, Lee JT, Beniwal S, Elezi G, et al. Discovery of a small molecule secreted clusterin enhancer that improves memory in Alzheimer’s disease mice. npj Drug Discovery. Springer Science and Business Media LLC; 2025;2. 10.1038/s44386-025-00009-2

54. Mahoney R, Ochoa Thomas E, Ramirez P, Miller HE, Beckmann A, Zuniga G, et al. Pathogenic Tau Causes a Toxic Depletion of Nuclear Calcium. Cell Rep. Elsevier B.V.; 2020;32. 10.1016/j.celrep.2020.107900

55. Berlind JE, Lai JD, Lie C, Vicente J, Lam K, Guo S, et al. KCTD20 suppression mitigates excitotoxicity in tauopathy patient organoids. Neuron. Cell Press; 2025;113:1169–1189.e7. 10.1016/j.neuron.2025.02.001

56. Huang H, Zheng S, Lu M. Downregulation of lncRNA MEG3 is involved in Parkinson’s disease. Metab Brain Dis. Springer; 2021;36:2323–8. 10.1007/s11011-021-00835-z

57. Yan H, Rao J, Yuan J, Gao L, Huang W, Zhao L, et al. Long non-coding RNA MEG3 functions as a competing endogenous RNA to regulate ischemic neuronal death by targeting miR-21/PDCD4 signaling pathway. Cell Death Dis. Nature Publishing Group; 2017;8. 10.1038/s41419-017-0047-y

58. Yi J, Chen B, Yao X, Lei Y, Ou F, Huang F. Upregulation of the lncRNA MEG3 improves cognitive impairment, alleviates neuronal damage, and inhibits activation of astrocytes in hippocampus tissues in Alzheimer’s disease through inactivating the PI3K/Akt signaling pathway. J Cell Biochem. Wiley-Liss Inc.; 2019;120:18053–65. 10.1002/jcb.29108

59. Yoon M, Kim H, Shin H, Lee HY, Kang MJ, Park SH, et al. Inhibition of CXXC5 function rescues Alzheimer’s disease phenotypes by restoring Wnt/β-catenin signaling pathway. Pharmacol Res. Academic Press; 2023;194. 10.1016/j.phrs.2023.106836

60. Solas M, Van Dam D, Janssens J, Ocariz U, Vermeiren Y, De Deyn PP, et al. 5-HT7 receptors in Alzheimer’s disease. Neurochem Int. Elsevier Ltd; 2021;150. 10.1016/j.neuint.2021.105185

