## Extended Data Figures for "Molecular Disease Stages of Oligodendrocytic and Neuronal Tau Burden in Progressive Supranuclear Palsy"


### Contributors and Institutions

Nils Briel[1,2,3], Viktoria C. Ruf[1], Paul Feyen[1,4], Sigrun Roeber[1], Thomas Arzberger[1], Otto Windl[1], Tobias Weiss[2], Paolo Arosio[3], Günter U. Höglinger[4,5,6,7], Felix L. Struebing[1,4], Jochen Herms[1,4]

1. Institute of Neuropathology, LMU Medizin, Ludwig-Maximilians-Universität (LMU) München, Munich, Germany
2. Department of Neurology, Zurich Neuroscience Center, University Hospital and University of Zurich, Switzerland
3. Department of Chemistry and Applied Biosciences, Swiss Federal Institute of Technology Zurich, Switzerland
4. German Center for Neurodegenerative Diseases, Site Munich, Germany
5. Department of Neurology, LMU University Hospital, LMU Medizin, Ludwig-Maximilians-Universität (LMU) München, Munich, Germany
6. Munich Cluster for Systems Neurology (SyNergy), Munich, Germany
7. Aligning Science Across Parkinson's (ASAP) Collaborative Research Network, Chevy Chase, MD, USA.

**Correspondence**:

Nils Briel, MD

Institute of Neuropathology, LMU Medizin

Ludwig-Maximilians-University Munich, Germany

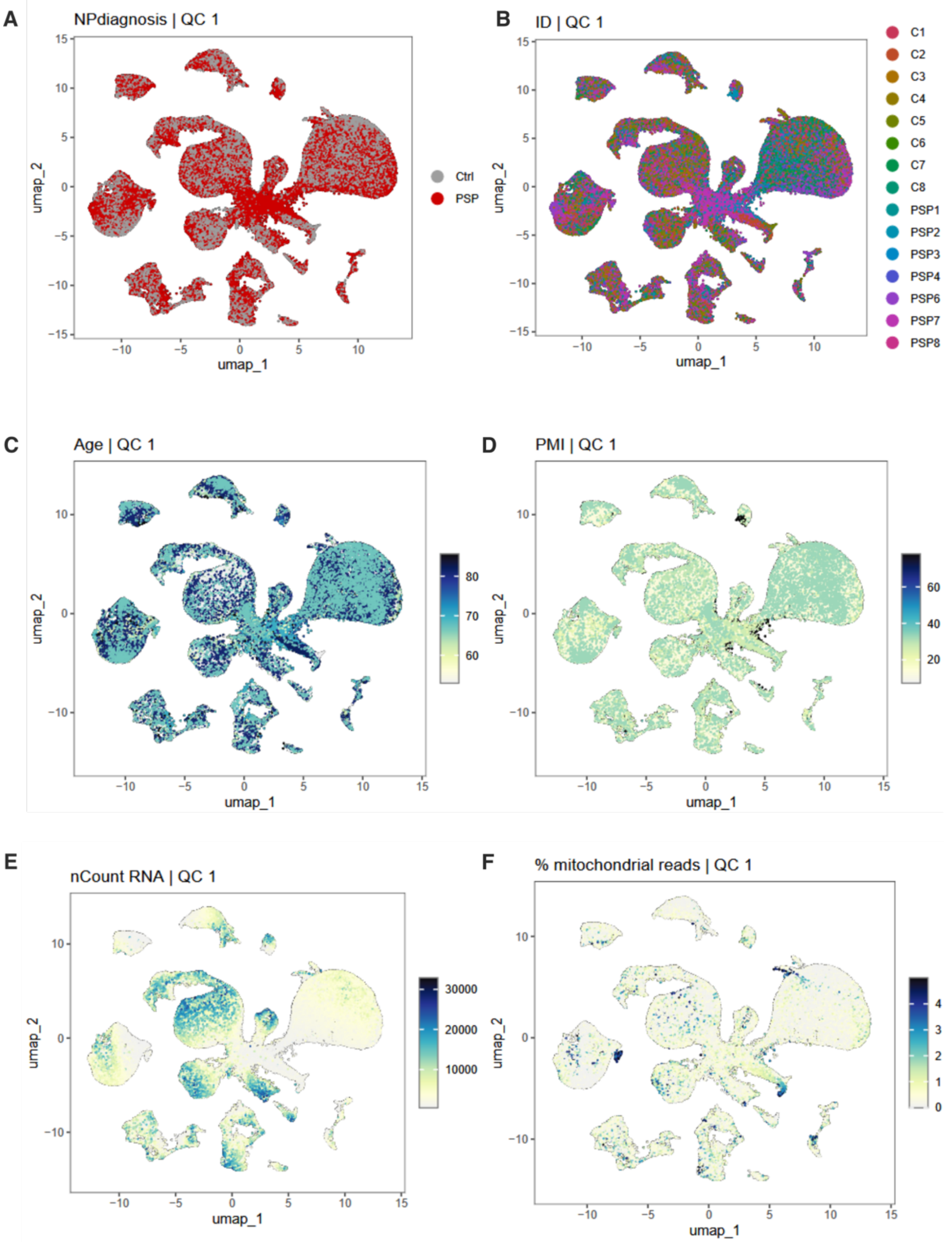


**<u>Extended Data Figure 1. snRNAseq QC 1 UMAP plots.</u>**

**(a)** UMAP of 60,038 nuclei colored by neuropathological diagnosis shows PSP (red) and control (grey) nuclei broadly intermingled, indicating limited disease-driven segregation after QC1.
**(b)** Identical UMAP colored by donor ID (14 brains) demonstrates uniform sample mixing and negligible batch structure.
**(c)** Nuclei colored by donor age (years) reveal an even age gradient across the manifold, with no age-specific clustering.
**(d)** UMAP colored by postmortem interval (PMI, hours) likewise shows no concentration of long-PMI nuclei.
**(e)** Library complexity per nucleus (nCount_RNA) plotted on UMAP highlights higher-complexity neuronal islands and lower-complexity glial zones.
**(f)** Percentage of mitochondrial reads (%MT) identifies sparse high-MT outliers that were flagged for removal in downstream QC.
<u>Abbreviations</u>: control (Ctrl, C1-8), RNA feature count (nCount RNA), neuropathological diagnosis (NPdiagnosis), postmortem interval (PMI).

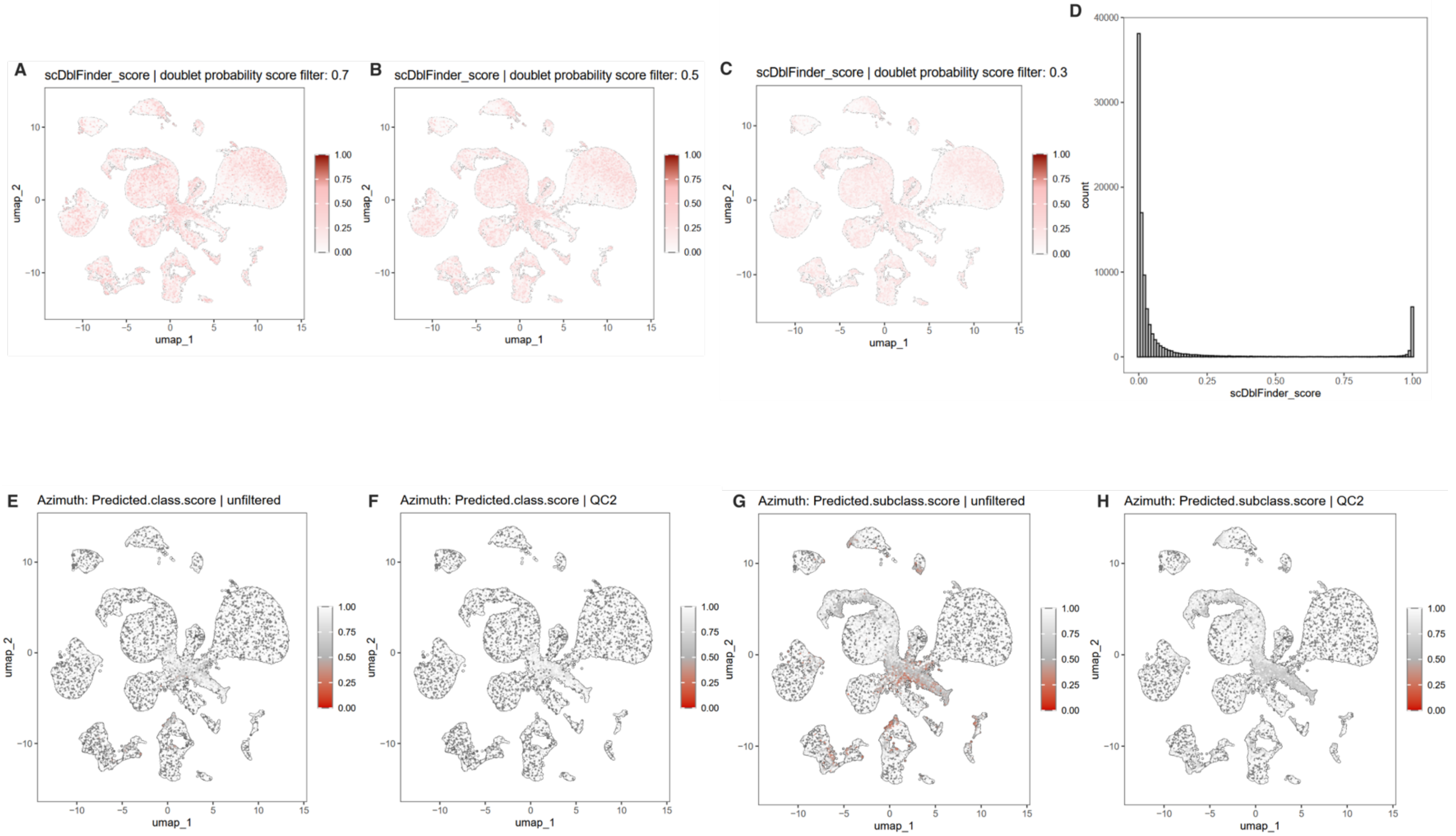


**<u>Extended Data Figure 2. snRNAseq doublet removal & cell identity labelling.</u>**

**(a-c)** UMAP projections colored by scDblFinder doublet probability illustrate nuclei flagged as potential doublets under increasingly stringent score cut-offs (0.7, 0.5, 0.3). Progressive filtering preferentially removes high-score nuclei while preserving cluster structure.
**(d)** Histogram of scDblFinder scores across all nuclei shows a right-skewed distribution, with the chosen cut-offs (vertical lines) separating a minority of high-probability doublets from the main singlet population.
**(e,f)** Azimuth predicted class scores mapped onto the UMAP before filtering (unfiltered) and after full QC (QC2). Post-QC nuclei exhibit uniformly high confidence (scores ≈ 1.0), confirming successful removal of ambiguous cells.
**(g,h)** Equivalent UMAPs for Azimuth predicted subclass scores demonstrate the same improvement in annotation confidence following QC2, with low-score nuclei largely absent after filtering.
<u>Abbreviations</u>: scDblFinder (single cell doublet finder, package name).

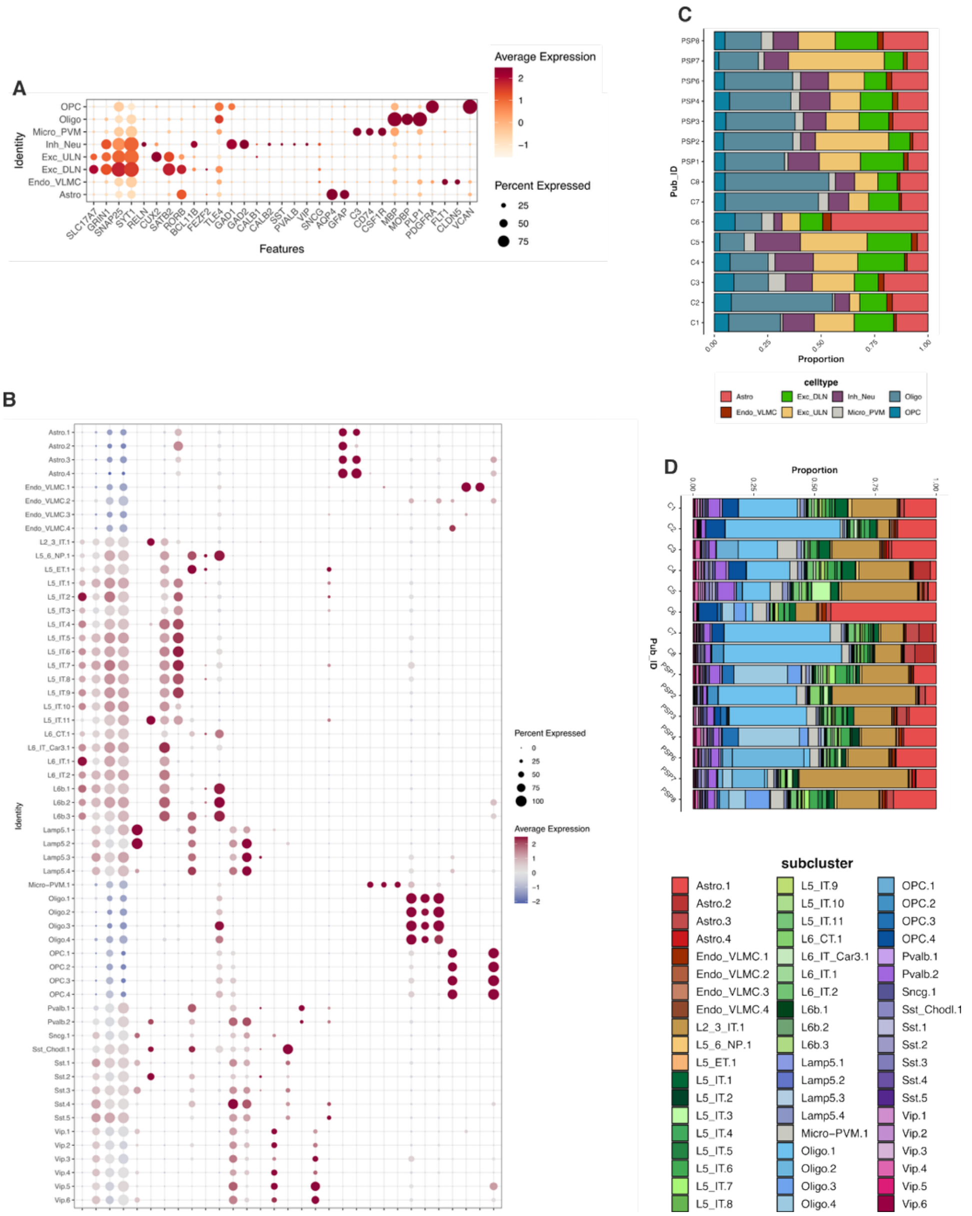


**Extended Data Figure 3. snRNAseq cell type marker gene expression.**

**(a)** Dot plot of canonical marker genes across **major cell classes**. Dot diameter encodes the percentage of nuclei expressing each gene, while color intensity reflects scaled average expression, highlighting clear class-specific signatures (e.g., Snap25 for neurons, Plp1 for oligodendrocytes).
**(b)** Higher-resolution dot plot displaying **subcluster-specific marker genes** within each major class, resolving laminar excitatory neuron subsets, interneuron subclasses, and glial subtypes; dot and color scales as in (a).
**(c)** Stacked bar charts summarizing **relative abundances of major cell classes** across individual samples, ordered by pathology group (P01–P09). Bars are partitioned by color-coded cell classes, facilitating visual comparison of broad compositional changes.
**(d)** Equivalent stacked bar charts for **fine-grained subclusters**, revealing nuanced shifts in laminar excitatory, inhibitory, and glial subpopulations across the same samples; color key lists all subcluster labels.
Abbreviations: control (Ctrl, C1-8), astrocytes (Astro), endothelial cells and vascular leptomeningeal cells (Endo_VLMC), excitatory deep-layer neurons (Exc_DLN), excitatory upper-layer neurons (Exc_ULN), inhibitory neurons (Inh_Neu), layer 2/3 intratelencephalic neurons (L2/3 IT), layer 5 extratelencephalic neurons (L5 ET), layer 5 intratelencephalic neurons (L5 IT), layer 5/6 near-projecting neurons (L5/6 NP), layer 6 corticothalamic neurons (L6 CT), layer 6 intratelencephalic neurons (L6 IT), microglia and perivascular macrophages (Micro_PVM), oligodendrocytic precursor cells (OPCs), oligodendrocytes (Oligo), parvalbumin-expressing interneurons (Pvalb), somatostatin-expressing interneurons (Sst), somatostatin/Chodl-expressing interneurons (Sst_Chodl), vasoactive intestinal peptide-expressing interneurons (Vip).

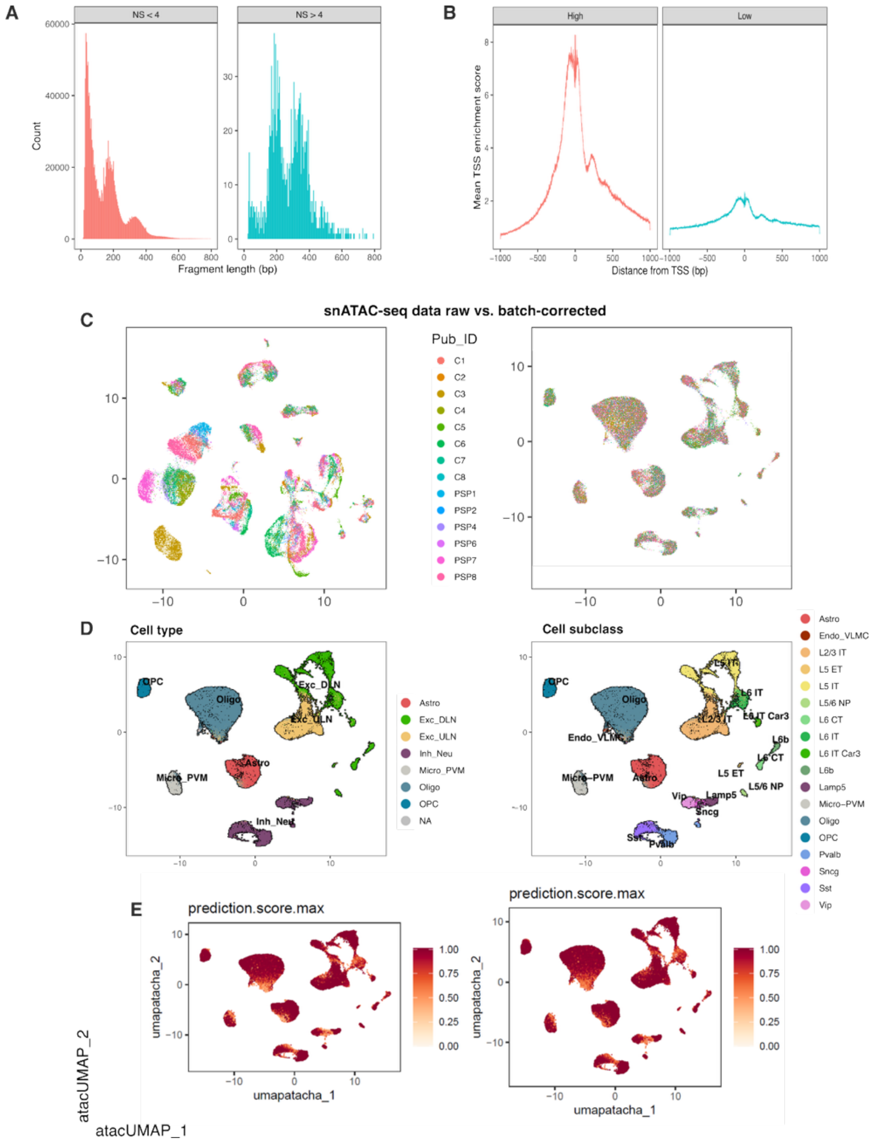


**Extended Data Figure 4. snATACseq QC.**

**(a)** Fragment length distribution histograms showing nucleosome-free regions and mono-, di-, and tri-nucleosomal peaks across samples, with insert size distributions validating chromatin accessibility profiling quality.
**(b)** TSS enrichment profiles demonstrating signal enrichment around transcription start sites, with mean TSS enrichment scores plotted against distance from TSS (±1kb), confirming successful ATACseq library preparation.
**(c)** snATACseq data raw vs. batch-corrected UMAP projections colored by publication ID, illustrating effective batch correction across different sample processing batches while preserving biological signal structure.
**(d)** Cell type and cell subclass annotations projected onto UMAP coordinates, showing distinct clustering patterns for major brain cell populations including astrocytes, excitatory neurons, inhibitory neurons, microglia, oligodendrocytes, and their respective subclusters.
**(e)** Prediction score maximum visualizations on atacUMAP coordinates, with color scales representing confidence scores for automated cell type classification, validating annotation accuracy across the single-nucleus chromatin accessibility landscape.
Abbreviations: base pairs (bp), control (Ctrl, C1-8), astrocytes (Astro), endothelial cells and vascular leptomeningeal cells (Endo_VLMC), excitatory deep-layer neurons (Exc_DLN), excitatory upper-layer neurons (Exc_ULN), inhibitory neurons (Inh_Neu), layer 2/3 intratelencephalic neurons (L2/3 IT), layer 5 extratelencephalic neurons (L5 ET), layer 5 intratelencephalic neurons (L5 IT), layer 5/6 near-projecting neurons (L5/6 NP), layer 6 corticothalamic neurons (L6 CT), layer 6 intratelencephalic neurons (L6 IT), microglia and perivascular macrophages (Micro_PVM), nucleosome signal (NS), oligodendrocytic precursor cells (OPCs), oligodendrocytes (Oligo), parvalbumin-expressing interneurons (Pvalb), somatostatin-expressing interneurons (Sst), somatostatin/Chodl-expressing interneurons (Sst_Chodl), transcription starting site (TSS), vasoactive intestinal peptide-expressing interneurons (Vip).

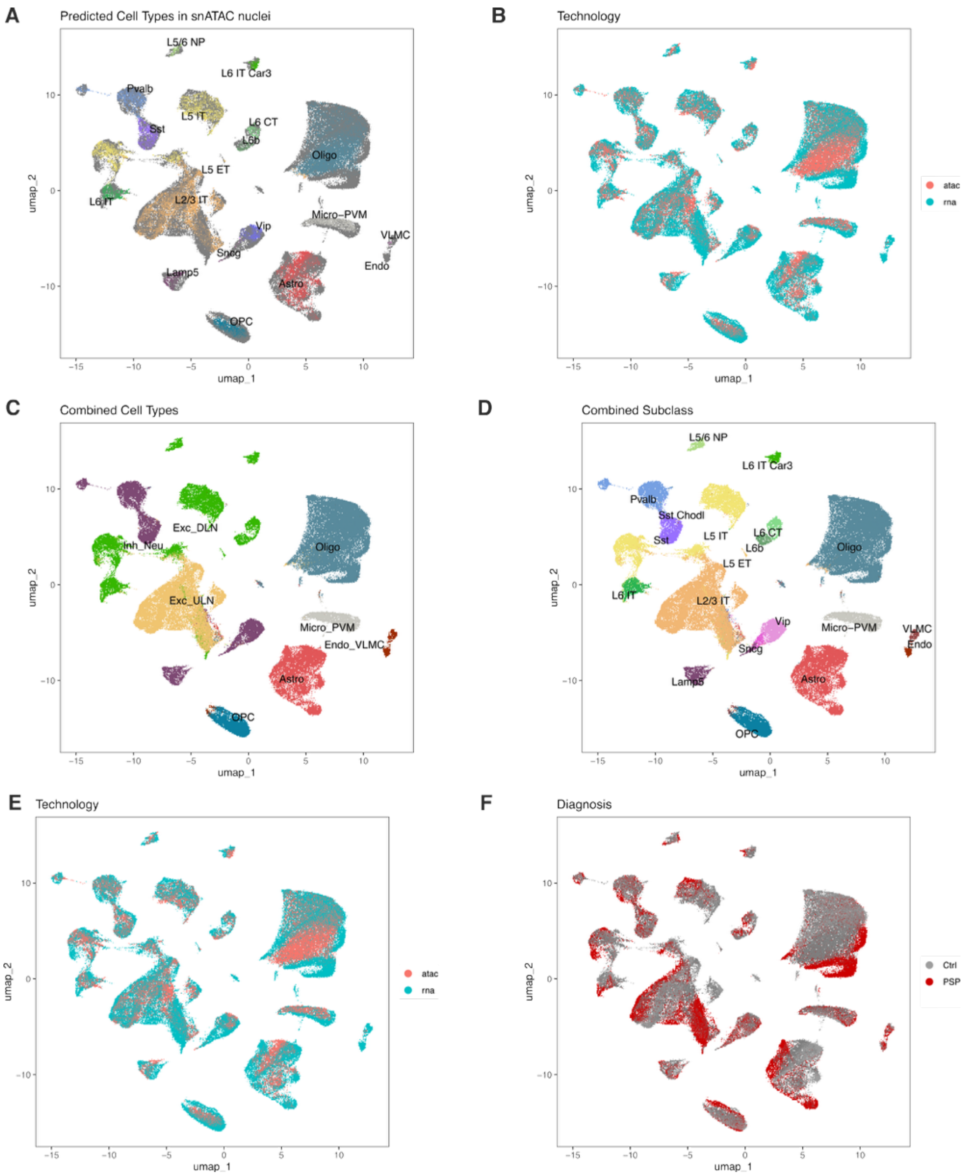


**Extended Data Figure 5. Coembedded snATAC/RNAseq UMAP plots.**

**(a)** UMAP of 64,305 snATAC nuclei after label, colored by predicted transcriptomic cell type, delineates excitatory neurons, inhibitory neurons, oligodendrocytes, astrocytes, microglia, endothelial cells, and VLMCs. **(b)** Identical UMAP colored by sequencing modality (ATAC vs RNA) shows tight intermixing of datasets, confirming successful cross-platform integration.
**(c)** Joint UMAP of the merged snATAC-snRNAseq dataset colored by consolidated cell type labels demonstrates concordant clustering across modalities.
**(d)** Higher-resolution view colored by subclass annotations resolves laminar excitatory subclasses (L2/3 IT, L4 IT, L5 IT, L6 IT, L6b) alongside major inhibitory and non-neuronal subclasses, supporting fine-grained label consistency. **(e)** Second integration replicate, colored by modality, recapitulates robust co-embedding and minimal batch structure. **(f)** Cells colored by diagnostic group (control, PSP, not assigned) distribute evenly across the manifold, indicating negligible diagnosis-driven segregation prior to downstream differential analyses.
Abbreviations: control (Ctrl), astrocytes (Astro), endothelial cells and vascular leptomeningeal cells (Endo_VLMC), excitatory deep-layer neurons (Exc_DLN), excitatory upper-layer neurons (Exc_ULN), inhibitory neurons (Inh_Neu), layer 2/3 intratelencephalic neurons (L2/3 IT), layer 5 extratelencephalic neurons (L5 ET), layer 5 intratelencephalic neurons (L5 IT), layer 5/6 near-projecting neurons (L5/6 NP), layer 6 corticothalamic neurons (L6 CT), layer 6 intratelencephalic neurons (L6 IT), microglia and perivascular macrophages (Micro_PVM), oligodendrocytic precursor cells (OPCs), oligodendrocytes (Oligo), parvalbumin-expressing interneurons (Pvalb), somatostatin-expressing interneurons (Sst), somatostatin/Chodl-expressing interneurons (Sst_Chodl), vasoactive intestinal peptide-expressing interneurons (Vip).

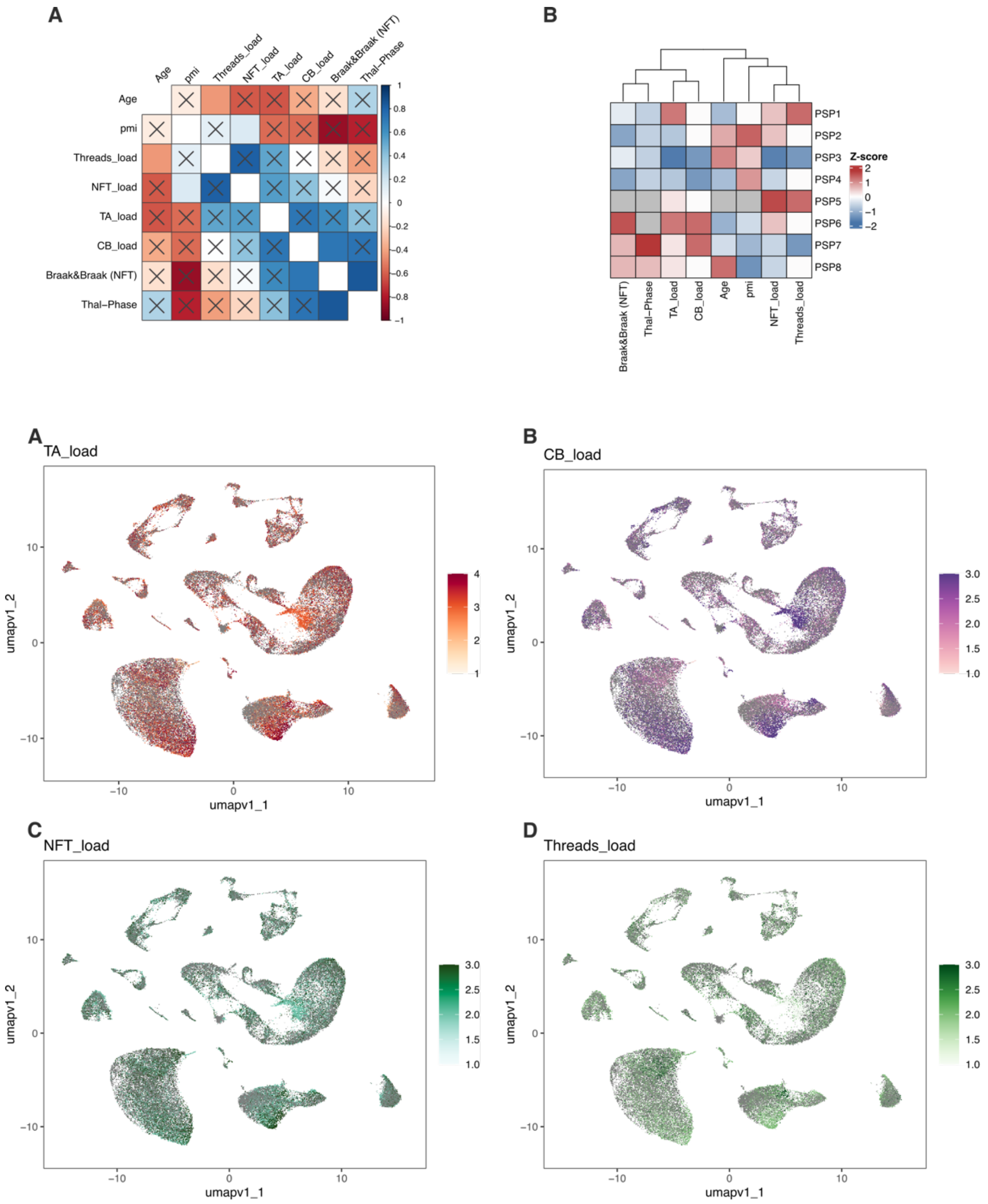


**Extended Data Figure 6. snRNAseq projection of cytopathological quantification**

**(a)** Correlation matrix showing pairwise relationships between age, PMI, and semi-quantitative cytopathological scores (TA, CB, NFT, Threads load; Braak/Break NFT stage; Thal-Phase) across all cases. Color scale encodes Pearson correlation coefficients; significant associations are marked with crosses.

**(b)** Hierarchical clustering heatmap of z-scored histopathological and clinical features across individual cases (PSP1–PSP7). Dendrograms display sample clustering based on shared quantitative profiles.

**(c)** UMAP projection of single-nucleus transcriptomes, colored by TA load, visualizes the spatial distribution of transcriptomic signatures corresponding to TA cytopathology severity.

**(d)** UMAP projection colored by CB load highlights the localization of transcriptomic patterns associated with CB cytopathology. Color scales in (c) and (d) represent semi-quantitative cytopathological scores.

NFT and Threads load are similarly projected on UMAP coordinates (not shown), with scales depicting respective pathology scores. All projections contextualize the relationship between cytoarchitectural heterogeneity and histological quantification in PSP.

<u>Abbreviations</u>: coiled bodies load (CB load), neurofibrillary tangle load (NFT load), postmortem interval (pmi), tufted astrocytes load (TA load).

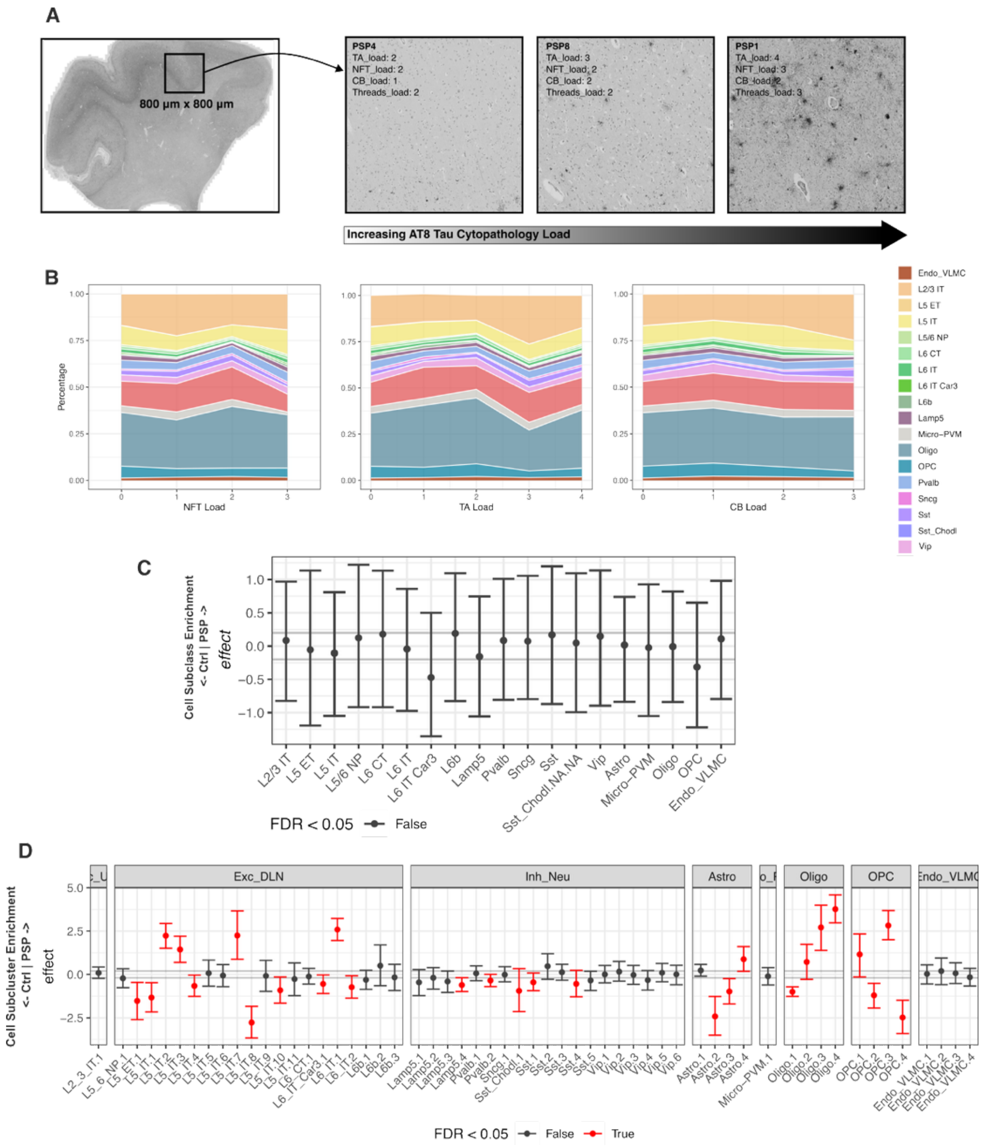


**Extended Data Figure 7. Cytopathological grading and cell type composition changes.**

**(a)** Representative tau immunostaining images from postmortem brain tissue illustrate a grading spectrum of AT8 tau cytopathology load across PSP cases, ranging from low to high severity at 800µm × 800µm scale.
**(b)** Stacked area plots depict proportions of major annotated cell populations across increasing neuropathological tau burden, assessed by NFT, TA, and CB load. Samples were stratified by semi-quantitative cytopathological scores, and color codes denote distinct cell types.
**(c)** Cell subclass enrichment effects estimated by *sccomp*, shown as one-dimensional credible intervals for each annotated population. Dots denote effect size directions and error bars uncertainty. Cell groups with significant enrichment or depletion (FDR<0.05) are highlighted per color scheme; markers at ±0.2 indicate effect size reference points.
**(d)** Cell subcluster enrichment effects estimated by *sccomp*, illustrating. Credible intervals and color coding as in (C), with significant differences marked (FDR<0.05).
Abbreviations: coiled bodies load (CB load), control (Ctrl), astrocytes (Astro), endothelial cells and vascular leptomeningeal cells (Endo_VLMC), excitatory deep-layer neurons (Exc_DLN), excitatory upper-layer neurons (Exc_ULN), false discovery rate (FDR), inhibitory neurons (Inh_Neu), layer 2/3 intratelencephalic neurons (L2/3 IT), layer 5 extratelencephalic neurons (L5 ET), layer 5 intratelencephalic neurons (L5 IT), layer 5/6 near-projecting neurons (L5/6 NP), layer 6 corticothalamic neurons (L6 CT), layer 6 intratelencephalic neurons (L6 IT), microglia and perivascular macrophages (Micro_PVM), neurofibrillary tangle load (NFT load), oligodendrocytic precursor cells (OPCs), oligodendrocytes (Oligo), parvalbumin-expressing interneurons (Pvalb), somatostatin-expressing interneurons (Sst), somatostatin/Chodl-expressing interneurons (Sst_Chodl), tufted astrocytes load (TA load), vasoactive intestinal peptide-expressing interneurons (Vip).

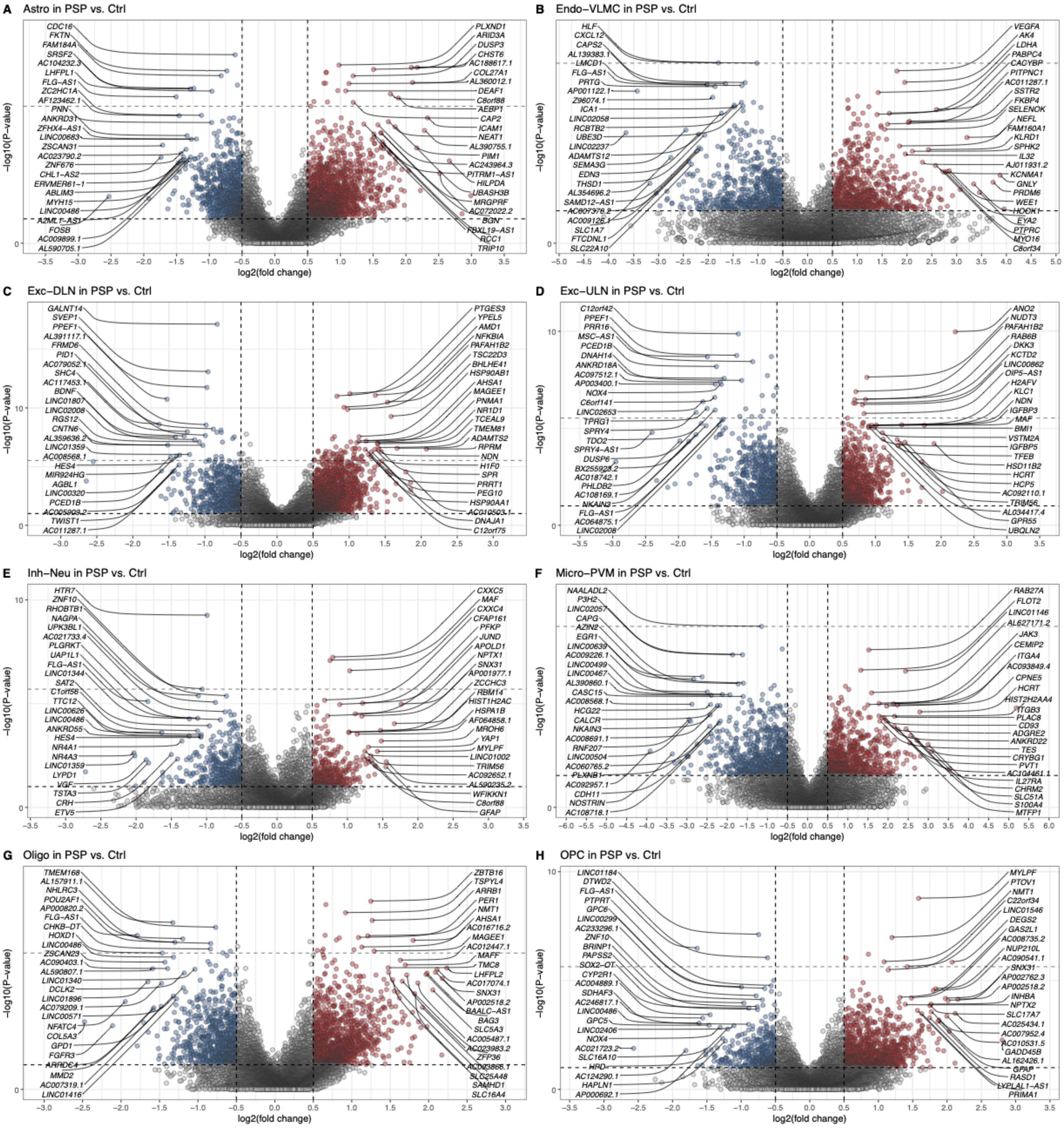


**Extended Data Figure 8. snRNAseq DEG analysis volcano plots.**

**(a-h)** Volcano plots show $log_2$FC vs. -$log_{10}$ p-value of DEGs for eight cell types in PSP (n = 7) versus controls (n = 8). Red and blue dots denote up- and downregulated genes (p-value < 0.05, |$log_2$FC| > 0.5), respectively; gray indicates non-significant or not fulfilling the $log_2$FC cutoff. Top 50 DEGs by FDR -$log_2$FC product are labeled. Vertical dashed lines mark ± 0.5 $log_2$FC; horizontal dashed lines mark p-value = 0.05 and FDR cutoff.

Abbreviations: astrocytes (Astro), control (Ctrl), differentially expressed genes (DEGs), endothelial cells and vascular leptomeningeal cells (Endo-VLMC), excitatory deep-layer neurons (Exc-DLN), excitatory upper-layer neurons (Exc-ULN), inhibitory neurons (Inh-Neu), $log_2$-fold-change ($log_2$FC), microglia and perivascular macrophages (Micro-PVM), oligodendrocytic precursor cells (OPCs), oligodendrocytes (Oligo).

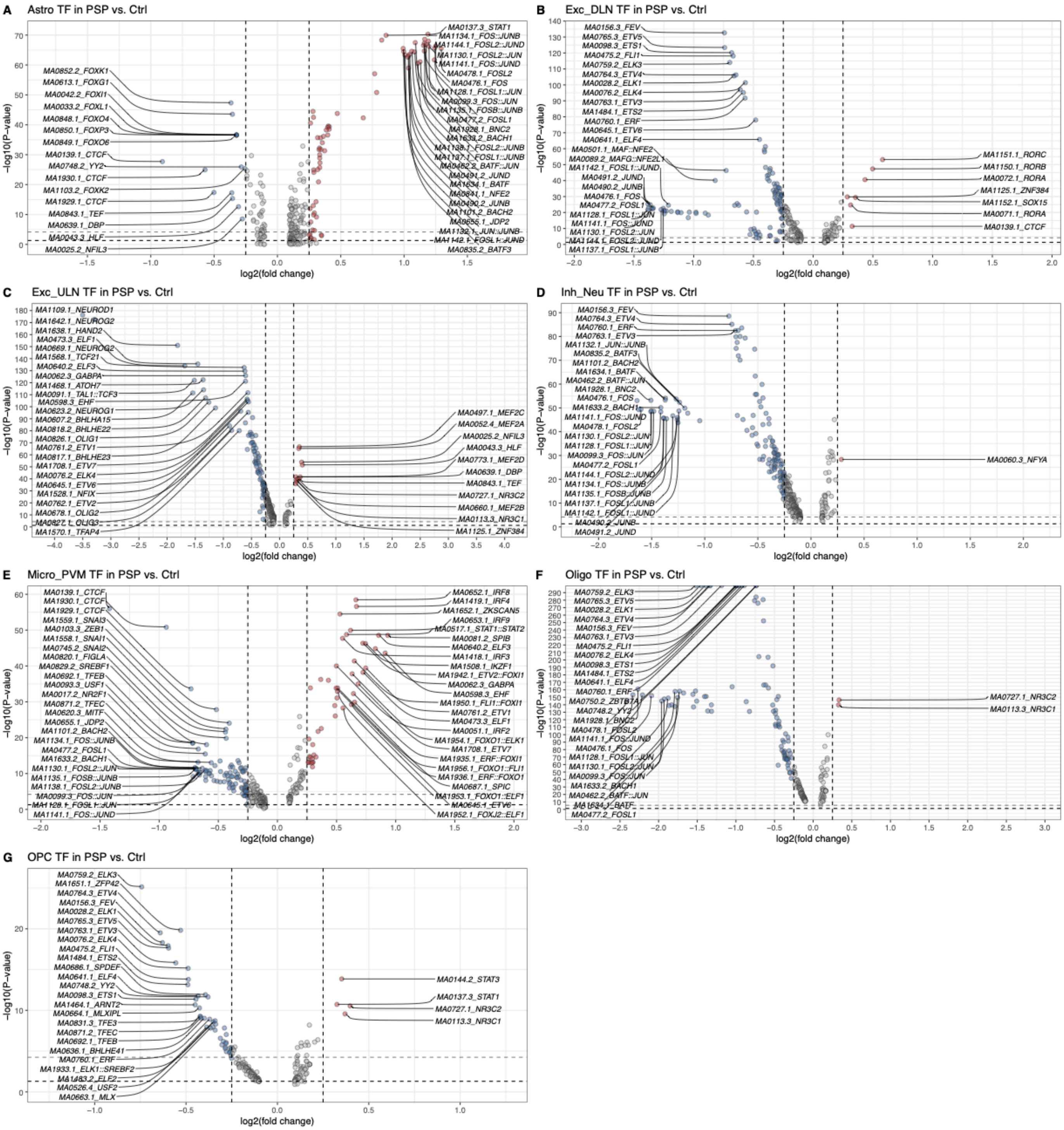


**Extended Data Figure 9. snATACseq DTF analysis volcano plots.**

**(a-h)** Volcano plots show $\log_2$FC vs. $-\log_{10}$ p-value of differentially regulated transcription factors (TF) for seven cell types (no significant findings in Endo-VLMC) in PSP (n = 6) versus controls (n = 8). Red and blue dots denote up- and downregulated TF motifs (p-value < 0.05, $|\log_2FC| > 0.25$); gray indicates non-significant or not fulfilling the $\log_2$FC cutoff. Top 50 TF by FDR $-\log_2$FC product are labeled. Vertical dashed lines mark ± 0.25 fold-change; horizontal dashed lines mark p-value = 0.05 and the FDR cutoff.

Abbreviations: astrocytes (Astro), control (Ctrl), endothelial cells and vascular leptomeningeal cells (Endo-VLMC), excitatory deep-layer neurons (Exc-DLN), excitatory upper-layer neurons (Exc-ULN), inhibitory neurons (Inh-Neu), $\log_2$-fold-change ($\log_2$FC), microglia and perivascular macrophages (Micro-PVM), oligodendrocytic precursor cells (OPCs), oligodendrocytes (Oligo), transcription factor (TF).

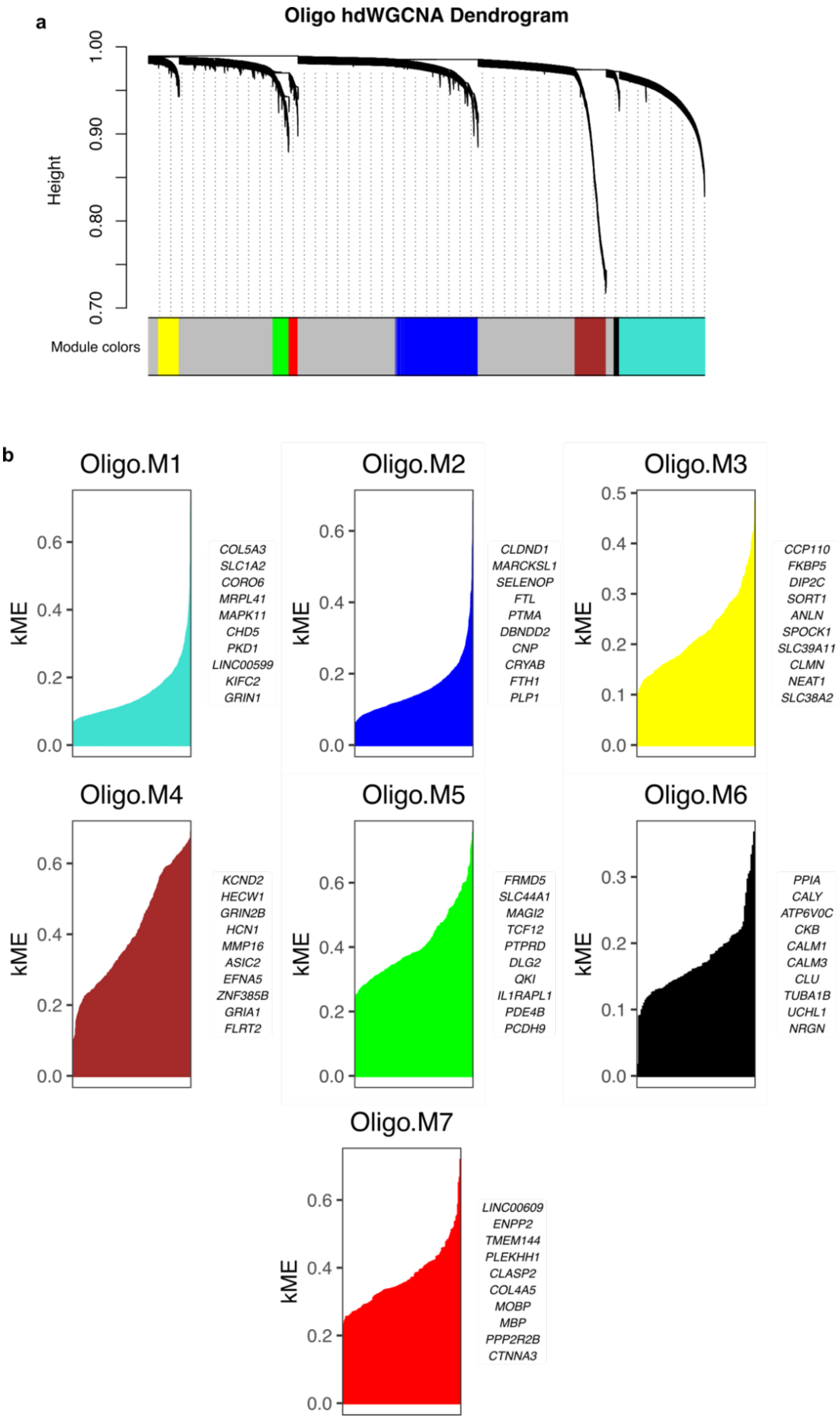


**Extended Data Figure 10. Characterization of oligodendrocytic co-expression networks via hdWGCNA.**

**(a)** Hierarchical clustering dendrogram showing gene modules (Oligo.M1-M7) based on topological overlap in the oligodendrocyte lineage. **(b)** Rank-ordered module membership (kME) plots for all co-expression modules. The top 10 hub genes by kME are listed for each module.
Abbreviations: high-dimensional weighted gene co-expression network analysis (hdWGCNA), oligodendrocytes (Oligo), k-module enrichment (kME)

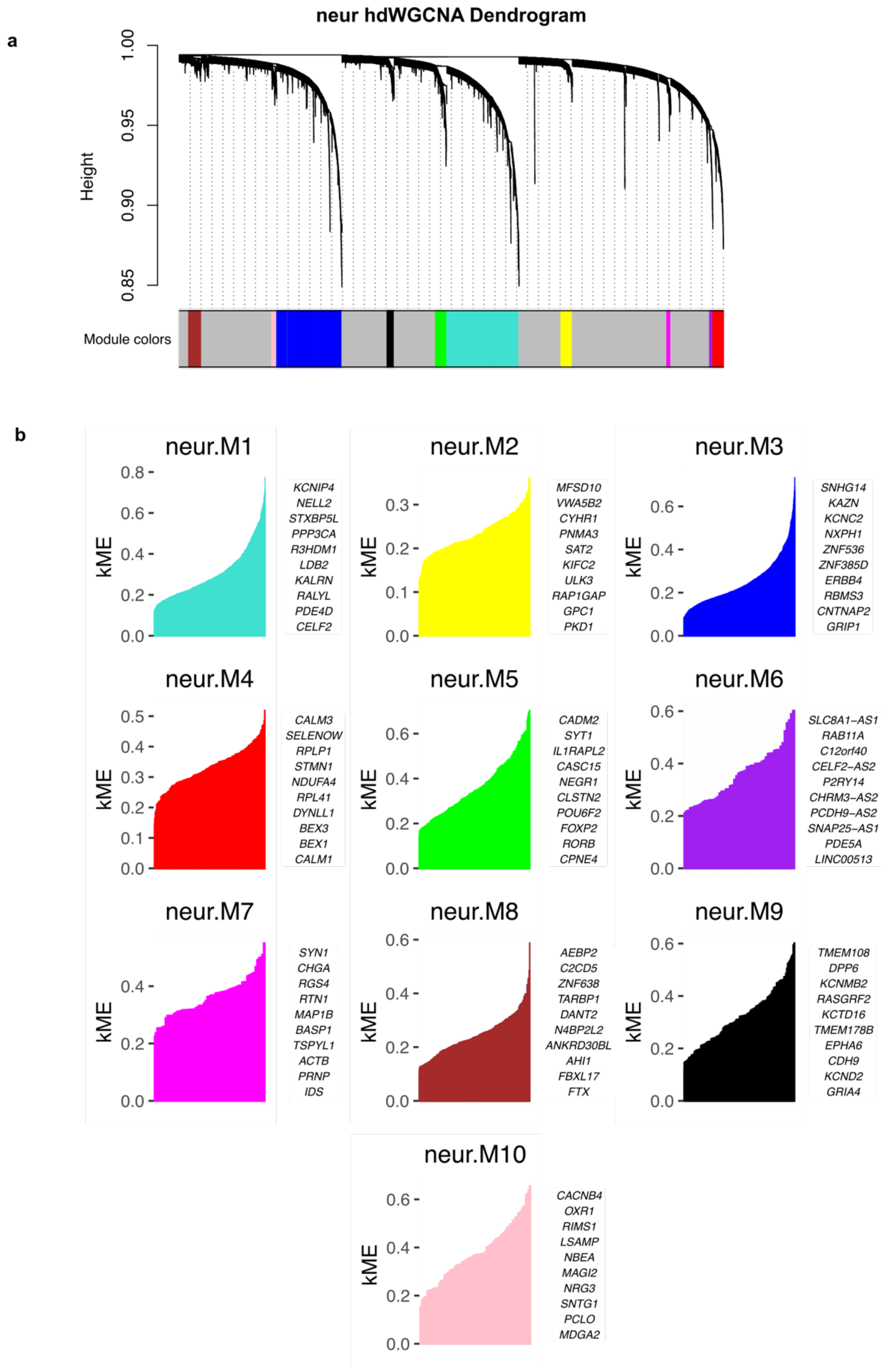


**Extended Data Figure 11. Characterization of neuronal co-expression networks via hdWGCNA.**

**(a)** Hierarchical clustering dendrogram showing gene modules (neur.M1-M10) based on topological overlap in the neuronal lineage. **(b)** Rank-ordered module membership (kME) plots for all co-expression modules. The top 10 hub genes by kME are listed for each module.

Abbreviations: high-dimensional weighted gene co-expression network analysis (hdWGCNA), pan-neurons (neur), k-module enrichment (kME)
